# COMBINATION THERAPIES TARGETING ALK-ABERRANT NEUROBLASTOMA IN PRECLINICAL MODELS

**DOI:** 10.1101/2022.10.30.512477

**Authors:** Elizabeth R. Tucker, Irene Jiménez, Lindi Chen, Angela Bellini, Chiara Gorrini, Elizabeth Calton, Qiong Gao, Harvey Che, Evon Poon, Yann Jamin, Barbara Martins da Costa, Karen Barker, Sumana Shrestha, J. Ciaran Hutchinson, Simran Dhariwal, Angharad Goodman, Elaine Del Nery, Pierre Gestraud, Jaydutt Bhalshankar, Yasmine Iddir, Elnaz Saberi-Ansari, Alexandra Saint-Charles, Birgit Geoerger, Maria Eugénia Marques Da Costa, Cécile Pierre-Eugène, Isabelle Janoueix-Lerosey, Didier Decaudin, Fariba Nemati, Angel M. Carcaboso, Didier Surdez, Olivier Delattre, Sally L. George, Louis Chesler, Deborah A. Tweddle, Gudrun Schleiermacher

**Affiliations:** Pediatric Tumour Biology and Therapeutics Team; Centre for Paediatric Oncology Experimental Medicine, Division of Clinical Studies; The Institute of Cancer Research, Cotswold Road, London, SM2 5NG, United Kingdom; SiRIC RTOP (Recherche Translationelle en Oncologie Pédiatrique); Translational Research Department, Institut Curie Research Center, PSL Research University, Institut Curie, Paris, France; INSERM U830, Equipe Labellisée Ligue contre le Cancer, PSL Research University, Institut Curie Research Center, Paris, France; Wolfson Childhood Cancer Research Centre, Translational & Clinical Research Institute, Newcastle Centre for Cancer, Newcastle University, Newcastle upon Tyne, United Kingdom; Division of Radiotherapy and Imaging; The Institute of Cancer Research, Cotswold Road, London SM2 5NG, United Kingdom; Newcastle Genetics Laboratory, Newcastle upon Tyne Hospitals NHS Trust, Newcastle upon Tyne, United Kingdom; Department of Histopathology, Great Ormond Street Hospital NHS Foundation Trust, London, UK; Department of Translational Research, The Biophenics High-Content Screening Laboratory, PSL Research University, PICT-IBiSa, Institut Curie Research Center, Paris, France; Bioinformatics platform, INSERM U900, PSL Research University, Institut Curie Research Center, Paris, France; Gustave Roussy Cancer Center, Department of Pediatric and Adolescent Oncology, Université Paris-Saclay, Villejuif, France; Laboratory of Preclinical Investigation, Department of Translational Research, Institut Curie, PSL University, Paris, France; Hospital Sant Joan de Déu, Barcelona, Spain; Balgrist University Hospital, Faculty of Medicine, University of Zurich (UZH), Zurich, Switzerland; Department of Medical Oncology, Institut Curie, Paris, France

**Keywords:** Neuroblastoma, ALK, GEMM, PDX, Idasanutlin

## Abstract

**Background:** *ALK* activating mutations are identified in approximately 10% of newly diagnosed neuroblastomas and *ALK* amplifications in a further 1-2% of cases. Lorlatinib, a third generation ALK inhibitor, will soon be given alongside induction chemotherapy for children with ALK-aberrant neuroblastoma. However, resistance to single agent treatment has been reported and therapies that improve the response duration are urgently required. We studied the preclinical combination of lorlatinib with chemotherapy, or with the MDM2 inhibitor, idasanutlin, as recent data has suggested that ALK inhibitor resistance can be overcome through activation of the p53-MDM2 pathway.

**Aims:** To study the preclinical activity of ALK inhibitors alone and in combination with chemotherapy or idasanutlin.

**Methods:** We compared different ALK inhibitors in preclinical models prior to evaluating lorlatinib in combination with chemotherapy or idasanutlin. We developed a triple chemotherapy (CAV: cyclophosphamide, doxorubicin and vincristine) *in vivo* dosing schedule and applied this to both neuroblastoma genetically engineered mouse models (GEMM) and patient derived xenografts (PDX).

**Results:** Lorlatinib in combination with chemotherapy was synergistic in immunocompetent neuroblastoma GEMM. Significant growth inhibition in response to lorlatinib was only observed in the *ALK*-amplified PDX model with the highest ALK expression. In this PDX lorlatinib combined with idasanutlin resulted in complete tumor regression and significantly delayed tumor regrowth.

**Conclusion:** Our study suggests that in neuroblastoma, high ALK expression could be associated with response to lorlatinib and either chemotherapy or idasanutlin. The synergy between MDM2 inhibition and ALK inhibition warrants further evaluation of this combination as a potential clinical approach for children with neuroblastoma.

**STATEMENT OF TRANSLATIONAL RELEVANCE:** Neuroblastoma is a pediatric tumor of the developing sympathetic nervous system. Around 50% of high-risk neuroblastoma patients are curable. Mutations or amplification of Anaplastic Lymphoma Kinase (ALK) have emerged as a marker with which to further risk-stratify patients. The ALK inhibitor lorlatinib will soon be used alongside chemotherapy in upfront treatment of high-risk patients with ALK-aberrant disease. In this preclinical study, we used a panel of *ALK* aberrant neuroblastoma models to evaluate ALK inhibitors focusing on lorlatinib in combination with conventional chemotherapy and the small molecule MDM2 inhibitor idasanutlin. In both approaches we found synergy in models with high basal ALK expression without MAPK pathway alterations. We conclude that in neuroblastoma the level of ALK expression could be an additional biomarker predictive of clinical response to ALK inhibitors.

## INTRODUCTION

Neuroblastoma is an embryonal tumor that arises in the developing sympathetic nervous system. Neuroblastoma is dramatically heterogeneous, ranging from spontaneous and complete regression (1) to very aggressive high-risk tumors. High-risk neuroblastoma defined as metastatic disease over one year of age or *MYCN* amplified disease, has a 5-year overall survival of 50% despite advances in treatment over the last 30 years (2) and accounts for approximately 15% of all childhood cancer deaths (3). New treatments and a better understanding of drug resistance are required to improve disease survival.

The heterogeneity of neuroblastoma is highly dependent on tumor biology. Whereas large scale copy number alterations occur in nearly all neuroblastoma as either numerical and/or segmental chromosome alterations, few recurrent molecular alterations targeting single genes have been described in neuroblastoma, of which some are correlated with poor outcome. *MYCN* amplification (4), *TERT* rearrangements (5–7), loss-of-function (LoF) *ATRX* mutations (8, 9) and *ALK* activating mutations and amplifications (10) are the most recognized. Both *MYCN* amplification and *TERT* re-arrangements lead to telomere maintenance by induction of telomerase, whereas *ATRX* LoF mutations induce telomere maintenance by activation of the alternative lengthening of telomeres (ALT) pathway. These alterations identify three almost non-overlapping groups of high-risk neuroblastomas, each associated with very poor prognosis, in particular when associated with mutations in the RAS/MAPK pathway (7–9, 11). As driver oncogenic events, these alterations could represent potential therapeutic targets for neuroblastoma.

*ALK* activating mutations are identified in approximately 10% of newly diagnosed neuroblastomas (12, 13) and *ALK* amplifications (14) in 1-2% of cases. Furthermore, the incidence of *ALK* mutations increases in relapsed neuroblastomas, occurring in around 20% of cases (15–17). Three mutation hotspots in the kinase domain (F1174, R1275 and F1245) represent 85% of all forms of *ALK* mutations (18). Together with *ALK* amplifications, these gain-of-function alterations lead to phosphorylation of ALK and constitutive increased kinase activity, as well as activation of downstream signaling molecules (such as PI3K-AKT, JAK-STAT and MAPK pathways), which results in enhanced cellular survival, migration and proliferation of neuroblastoma (12, 13). In addition, germline *ALK* mutations are the major cause of hereditary neuroblastoma, which account for 1-2% of all neuroblastoma cases (12, 13, 19). The high incidence of ALK activation justifies the study of ALK inhibition as a therapeutic target in neuroblastoma.

The most extensively studied ALK inhibitor in neuroblastoma, crizotinib, is a small molecule competitive inhibitor of ALK and MET kinase activity and was the first FDA approved drug for use in adult patients with *ALK*-translocated non-small cell lung cancer (NSCLC). A phase II trial involving crizotinib has been completed by the Children’s Oncology Group (COG) in pediatric patients with relapsed/refractory ALK-driven neuroblastoma (NCT00939770). Only 3 of 20 patients showed objective responses (20), mainly explained by the intrinsic resistance of F1174 and F1245 hotspot mutations to crizotinib (18). Preclinical studies showed that crizotinib resistance can be overcome when combined with chemotherapy (21), and was the rationale for the COG phase I trial combining crizotinib with topotecan and cyclophosphamide in children with relapsed and refractory solid tumors (NCT01606878) (22). The second generation ALK inhibitor ceritinib resulted in objective responses in patients with neuroblastoma harboring ALK F1275 mutations whereas patients with F1174 mutations did not benefit (23).

Later generation ALK inhibitors have been developed to overcome both intrinsic and *de novo* resistance observed with crizotinib and ceritinib. Lorlatinib, a third generation ROS1 and ALK inhibitor has been shown to exert preclinical activity against cell line xenografts, patient-derived xenografts (PDX) and genetically engineered mouse models (GEMM) harboring all hotspot ALK mutations (24, 25). These findings led to a phase I trial of lorlatinib for patients with *ALK*-driven neuroblastoma that is currently ongoing through the New Approaches to Neuroblastoma Therapy (NANT) consortium (NCT03107988), but even following initial response, relapse can occur. Resistance to ALK inhibition is thought to occur through several different mechanisms, including changes in the epigenetic landscape and re-activation of the RAS-MAPK pathway (26, 27).

Furthermore it has been shown that it is also possible to overcome resistance to ALK inhibition by activating the p53-MDM2 pathway (28, 29). In contrast to other malignancies, neuroblastomas rarely harbor *TP53* mutations. Nevertheless, there is evidence of p53 pathway inactivation in 15% of neuroblastomas at the time of relapse, that may contribute to chemotherapy resistance (30). Several mechanisms for p53 inactivation have been proposed, including MYCN–mediated MDM2 overexpression or *MDM2* gene amplification or homozygous p14 deletion (31–33). Thus, enhancing or reactivating the functional activity of p53 by targeting the p53-MDM2 pathway via MDM2 inhibition may represent a plausible approach for neuroblastoma treatment.

We hypothesized that combining lorlatinib with chemotherapy or MDM2 inhibitors will lead to enhanced efficacy in neuroblastoma and, ultimately in a clinical setting, improved patient survival. In this work, the *in vitro* and *in vivo* activity of ALK inhibitors has been explored alone and in combination with chemotherapy or MDM2 inhibitors in various *ALK*-mutated or amplified preclinical models of neuroblastoma, including *in vitro* (cell lines), *ex vivo* (PDX-derived tumor cells, PDTCs), and *in vivo* GEMM and PDX models. These models were also used to study the activity of the combination of ALK and MDM2 inhibition for the treatment of *ALK*-aberrant neuroblastomas. We found that in the context of ALK aberrant neuroblastoma, treatment efficacy with lorlatinib, either alone or as part of combination therapy, was associated with high basal ALK expression, and we suggest this might be evaluated as an additional biomarker in addition to genetic status in future neuroblastoma clinical trials of ALK inhibitors in neuroblastoma.

## MATERIALS AND METHODS

### Treatment of ALK aberrant neuroblastoma cell lines

Human neuroblastoma cell lines were cultured in RPMI 1640 (Sigma) supplemented with 10% v/v Fetal Calf Serum (FCS; Gibco/Life Technologies Ltd, Paisley, UK) (**Supplementary Table S1**) (30). Cell lines were authenticated using STR genotyping and/or cytogenetic analyses. Cell lines were sequenced for *ALK* by Sanger sequencing (exons 20-29) and targeted next generation sequencing for ALK and RAS-MAPK pathway genes (34) and *ALK* amplification was assessed by FISH and/or SNP array, the latter analyzed using Nexus Biodiscovery software (**Supplementary Fig. S1A**).

The MDM2 antagonist idasanutlin was provided by Hoffman-La Roche/Genentech. All other compounds were purchased from Selleck Chemicals (Stratech Scientific Ltd, Newmarket, UK). All compounds for *in vitro* studies were dissolved in dimethyl sulfoxide (DMSO) (Sigma-Aldrich). Seventy-two-hour XTT growth inhibition assays with determination of the concentration required to inhibit growth by 50% (GI_50_) and median effect analysis were performed as previously described in cells growing exponentially (35). Synergy and Combination Index (CI) values with the ALK inhibitors TAE684 and alectinib (as a prelude to *in vivo* studies with the clinical inhibitor lorlatinib) in combination with idasanutlin, were determined using CalcuSyn v2 (Biosoft, Cambridge, UK). Flow cytometry and caspase 3/7 assays were performed as previously described and according to manufacturer’s protocols (35).

### *Ex vivo* drug screenings in PDTC models

#### Obtaining PDTCs from PDX tumors

For viable PDTC generation, we used a dissociation protocol adapted from Stewart *et al*., 2017 (36), which included a first step of PDX tumor mechanical dissociation with sterile scalpels and then enzymatic dissociation by trypsin (10 mg/ml) and type II collagenase (275 U/mg, Worthington Biochemical). The tube was then placed in a warm 37°C water bath for 60 min. Dissociation was stopped by adding Soybean Trypsin Inhibitor (10 mg/ml, Sigma). Deoxyribonuclease I (2 mg/ml, Sigma) and magnesium chloride (1 M) were added in equal amounts. Tumor suspension was filtered with a 40 μm cell strainer and then centrifuged at 500g for 5 minutes. Supernatant was discarded and the cell pellet was resuspended in PBS-minus/10%FBS for cell counting. The suspension was then recentrifuged and resuspended in serum-free stem-cell (SC) medium, which contained Dulbecco’s Modified Eagle Medium/Nutrient Mixture F-12 (DMEM/F12, Gibco) supplemented with 40 ng/ml basic fibroblast growth factor (bFGF), 20 ng/ml epidermal growth factor (EGF), 1× B27 supplement (Gibco) and 500 U/ml of penicillin/streptomycin.

#### Experimental pipeline

The same experimental pipeline was used for all drug screens in PDTCs. All compounds for PDTC experiments were purchased from Selleck Chemicals (Stratech Scientific Ltd, Newmarket, UK) and dissolved in DMSO (Sigma-Aldrich). After PDX tumor dissociation, cells in SC medium were plated by robotic seeding in 384-well plates at a concentration of 20,000 cells in 40 μL per well. Drugs were added to cells 24 hours after plating (day 1) by a robotic drug dispenser. Cells were then incubated for 72 hours until day 4, when cell viability was measured by CellTiter Glo (Promega) luminescent assay.

#### Monotherapy drug screening

The same experimental pipeline was used for all drug screens in PDTCs. ALK inhibitors crizotinib, alectinib, ceritinib, and lorlatinib were purchased from Selleck Chemicals (Stratech Scientific Ltd, Newmarket, UK) as 10 mM stock solution in 100% DMSO. PDTCs in SC medium were seeded in 384-well plates (ViewPlate-384 Black Perkin Elmer, ref. 6007460) using a MultiDrop combi (Thermo Fisher Scientific) at a concentration of 20,000 cells in 40 μL per well. Twenty-four hours after cell seeding, drugs were robotically added to each well and titrated by three-fold dilutions covering 12 concentrations from 10,000nM to 0.056nM, across three 384-well plates. Control wells received the same distributed volumes with either DMSO (DMSO control) or cell media alone (empty). Plates were then incubated at 37°C in 5% CO_2_ for 72h, then cell viability was evaluated in culture based on quantification of the ATP present using CellTiter Glo Luminescent cell viability assay (Promega). Raw luminescence was recorded using a CLARIOStar (BMG Labtech) and obtained values were normalized on a per-plate basis by dividing the raw luminescence value in each well by the median value obtained for the DMSO-treated wells condition (100% viability). Dose-response curves were fitted with a 4 parameters log-logistic model with a fixed viability of 100% at 0nM using R package drc. Drug responses were represented by the quantitative drug sensitivity score (DSS), as described by Yadav *et al*. (37). Each PDX model was screened in two independent biological replicates (two different experiments for each PDX model, at a different PDX passage number, with high correlation across the results (**Supplementary Fig. S2A**)).

### *In vivo* experiments with GEMMs

All experiments, including the breeding of transgenic animals, were performed in accordance with the local ethical review panel, the UK Home Office Animals (Scientific procedures) Act 1986, the ARRIVE (Animal Research: Reporting of In Vivo Experiments) guidelines (38) and the UK NCRI guideline (39).

For GEMM experiments crizotinib and lorlatinib were provided by Pfizer, alectinib was provided by Hoffman-La Roche/Genentech, and ceritinib was purchased from Selleck Chemicals (Stratech Scientific Ltd, Newmarket, UK).

Th-*ALK*^F1174L^/*MYCN* tumor-bearing animals were enrolled into therapeutic trials when their abdominal tumors reached 5 mm in diameter according to palpation. Changes in tumor volume in the TH-*ALK^F1174L^/*TH-*MYCN* mice were quantified using MRI on a 7T horizontal bore MicroImaging system (Bruker Instruments) using a 3 cm birdcage coil. Anatomical T_2_-weighted coronal images were acquired through the mouse abdomen, from which tumor volumes were determined using segmentation from regions of interest (ROI) drawn on each tumor-containing slice (40).

#### Short-term studies

For *in vivo* oral dosing, crizotinib was dissolved in sterile water with 10% Tween 20. Ceritinib was dissolved into 0.5% methylcellulose, 0.5% Tween 80 with sterile water. Alectinib was dissolved in 10% DMSO, 10% cremophor, 15% PEG400, 15% HPCD, 0.02N HCl in sterile water. Lorlatinib was dissolved into 0.5% methylcellulose, 0.5% Tween 80 with sterile water. The following doses were used for CAV chemotherapy, given as two intraperitoneal injections: (C) cyclophosphamide: 28 mg/kg; (A) doxorubicin: 0.7 mg/kg; (V) vincristine: 0.015 mg/kg (41). Two hours following the final dose of compound, tumor tissue was excised and snap frozen prior to analysis.

#### Longer-term survival studies

For longer-term survival studies tumor volume was monitored by daily palpation, and the animal was sacrificed when UK Home Office License limits were reached to record survival. Daily dosing of compounds was continuous for maximum of 56 doses. Mice receiving the CAV chemotherapy regimen were treated with an intraperitoneal injection on day 1 with doses as described above. In the combination regimen, lorlatinib was commenced from day 2 at 10 mg/kg orally, once daily. In the single drug regimen, lorlatinib was commenced on day 1 at 10mg/kg orally, twice daily. Mice were sacrificed 2 hours following the final dose of lorlatinib and any remaining tumor tissue collected for pharmacodynamic studies.

#### Pharmacodynamic studies on GEMM tumors

Snap frozen tumors were lysed in 5% CHAPS buffer and quantitation of ALK and pY1586 ALK was performed using immunoassay as previously described (42).

### PDX *in vivo* experiments

All *in vivo* experiments on PDX were performed according to the European and National Regulation in force for the Protection of Vertebrate Animals used for Experimental and other Scientific Purposes (Directive 86/609). This study was approved by the Ministère de l’Education Nationale, de l’Enseignement Supérieur et de la Recherche (authorization number APAFIS # 13980-2018030813227748 v2).

#### PDX treatments

*Swiss Nude* mice (Charles River) were engrafted in the interscapular fat pad. Twelve mice per group were engrafted per experimental arm to ensure at least 5 treated mice and 3 PDX tumors for pharmacodynamic studies. For PDX experiments all drugs were purchased from Selleck Chemicals (Stratech Scientific Ltd, Newmarket, UK), and dissolved as per the GEMM experiments. Treatment started when tumor reached 60-180 mm^3^. All mice received the CAV chemotherapy regimen intraperitoneally (IP) on day 1 (D1) with the same doses as used in the GEMM. For efficacy studies, mice received idasanutlin and lorlatinib orally at 75 mg/kg/dose from D1 and 10 mg/kg/dose from day 2 (D2), respectively, 5 days per week. Targeted therapies were continued until tumor ethical size at 2,000 mm^3^. For pharmacodynamic studies, mice received CAV chemotherapy on D1, idasanutlin on D1 and D2, and lorlatinib on D2 and D3 (all drugs at the doses described above). Mice were sacrificed on D3, 4 hours after the second dose of lorlatinib, for pharmacodynamic studies.

#### Antitumor efficacy assessment on PDX tumors

Tumor volumes were calculated by measuring two perpendicular diameters with calipers. Each tumor volume (V) was calculated according to the following formula: V = a × b^2^ / 2, where a and b are the largest and smallest perpendicular tumor diameters. Relative tumor volumes (RTV) were calculated from the following formula: RTV = (Vx/V1), where Vx is the tumor volume on day x and V1 is the tumor volume at initiation of therapy (D1). Growth curves were obtained by plotting the mean values of RTV on the Y axis against time (X axis, expressed as days after the start of treatment). Antitumor activity was evaluated according to tumor growth inhibition (TGI), calculated according to the following formula: percent GI = (1 - RTVt / RTVc) × 100, where RTVt is the median RTV of treated mice and RTVc is the median RTV of controls, both at a given time point when the antitumor effect was optimal. Fifty percent TGI was considered to be the limit for a meaningful biological effect. Statistical significance of differences observed between the individual RTVs corresponding to the treated mice and control groups was calculated by the two-tailed Student’s t test.

#### Pharmacodynamic studies on PDX tumors

Tumors were extracted on D3 of treatment, 4 hours after the second dose of lorlatinib. All tissue blocks were cut to provide sections of 3□μm. Immunohistochemistry was performed for Ki67 (Abcam ab15580), cleaved caspase-3 (Cell Signaling #9661) and ALK to assess for cell proliferation, apoptosis, and the level of ALK expression respectively. The evaluation of immunostaining was conducted by two independent pathologists. The intensity of the immunostaining in the cell cytoplasm and the percentage of neoplastic cells with positive immunostaining were estimated. The score is the result of the multiplication of color intensity and percentage of cells with positive immunostaining, giving scores in the range of 0–3. Western blotting was carried out for ALK (Cell Signaling #3633), pY1586 ALK (Cell Signaling #3348), ERK1/2 (Cell Signaling #4695), pERK1/2 (Cell Signaling #9101), AKT (Cell Signaling #9272), pAKT (Cell Signaling 4060#) and GAPDH (Cell Signaling #2118) as previously described. (35, 43). Densitometry was performed using ImageJ.

## RESULTS

### Lorlatinib is the most potent ALK inhibitor in both *in vitro* and *in vivo* models of neuroblastoma

Given the poorer prognostic impact of *ALK* amplifications and clonal *ALK* mutations in high-risk neuroblastoma patients (10), we sought to further study the pre-clinical efficacy of ALK inhibitors in neuroblastoma models. In a first step, GI_50_ values for the ALK inhibitors TAE-684 (a preclinical ALK inhibitor included to provide comparison with previous studies), crizotinib, alectinib, ceritinib and lorlatinib were determined in a panel of neuroblastoma cell lines of varying *ALK* status using 72-hour XTT assays. Sensitivity to the tested ALK inhibitors except for lorlatinib correlated with the presence and type of *ALK* alteration (i.e., mutation or amplification) (**Fig. 1 Ai-v**). Cell lines with *ALK* alterations were generally less sensitive to lorlatinib than crizotinib, with a much wider range of reported GI_50_ values also seen for lorlatinib. A recent paper reported that the co-existence of RAS/MAPK pathway mutation following ALK inhibitor treatment confers resistance to ALK inhibitors, particularly the most selective ALK inhibitor lorlatinib, due to by-pass ALK signaling (27). Using targeted NGS we found that *ALK* aberrant cell lines most resistant to lorlatinib were those with additional RAS/MAPK pathway mutations e.g., NBLW, LAN6, SH-SY5Y (**Supplementary Table 1**).

**Figure 1.**
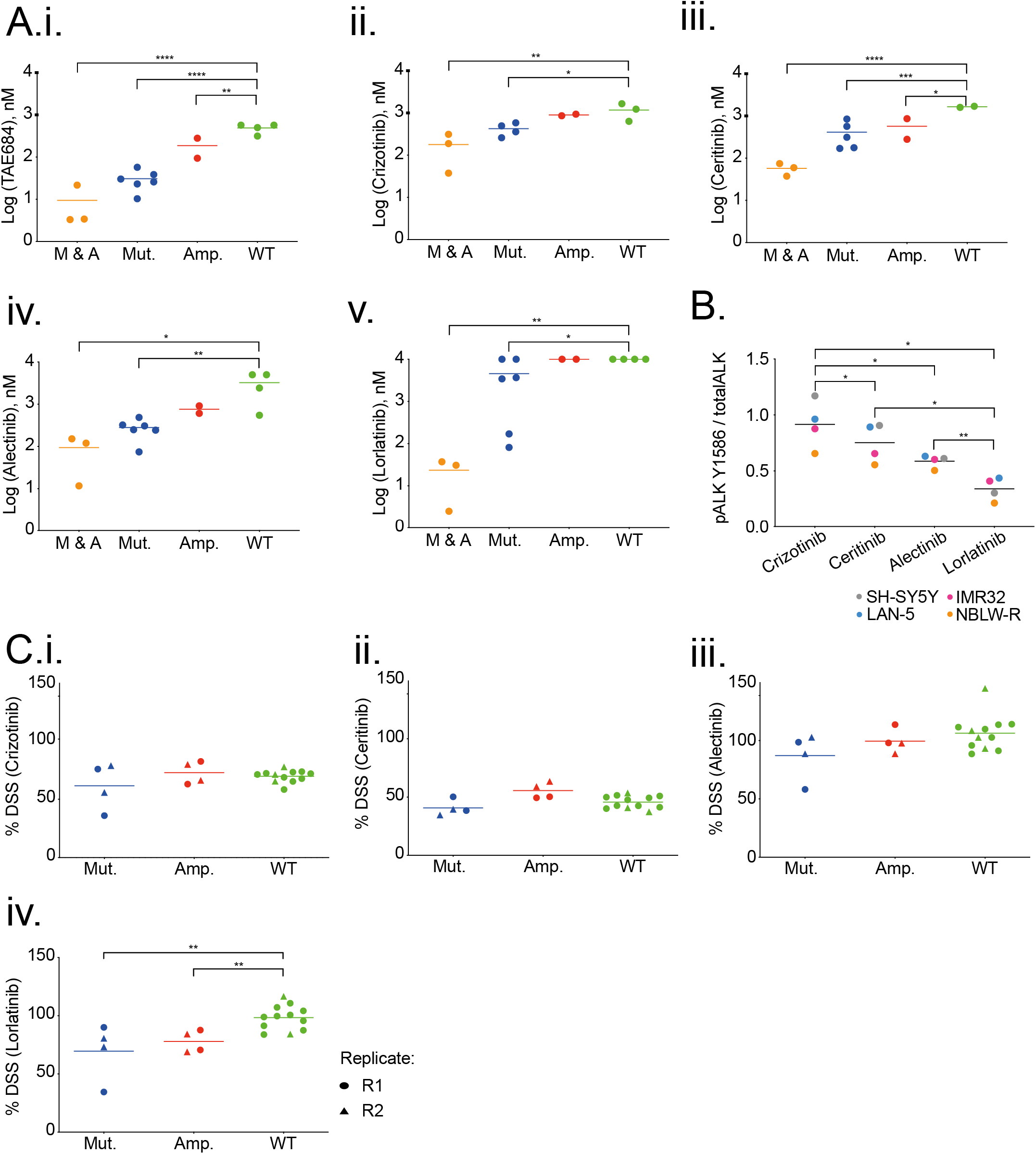
Lorlatinib is the most potent ALK inhibitor tested across *ALK*-mutant or *ALK*-amplified neuroblastoma cell lines and PDTC *ex vivo* models. 72-hour GI_50_ values for ALK inhibitors. (A), (i) TAE-684, (ii) crizotinib, (iii) ceritinib, (vi) alectinib and (v) lorlatinib in a panel of neuroblastoma cell lines (M & A: mutant and amplified *ALK*; Mut.: mutant *ALK*; Amp.: amplified *ALK*; WT: wild type *ALK*). Cell lines were grouped based on the type of *ALK* alteration. Statistically significant differences were determined by one-way ANOVA with Bonferroni post-hoc tests and paired testing versus WT. P ≤ 0.05 (*); 0.01 (**); 0.001 (***); 0.0001 (****). (B) NB cell lines treated with 20nM of indicated inhibitor for 3 hours, and lysates subjected to ALK immunoassay for total ALK and pY1586 ALK. Statistically significant differences determined by paired, two-tailed t-test. P≤0.05 (*); 0.01 (**); 0.001 (***). (C) Study of cell viability by analysis of the drug sensitivity score (DSS) after treatment with (i) crizotinib, (ii) ceritinib, (iii) alectinib or (iv) lorlatinib in *ALK*-mutant and amplified PDTC models. R1: replicate 1; R2 replicate 2. Statistically significant differences determined by unpaired, two-tailed t-test. P≤ 0.01 (**).

We quantified *in vitro* inhibition of ALK phosphorylation by immunoassay, comparing the different ALK inhibitors in a subset of our panel of neuroblastoma cell lines and found that lorlatinib was the most potent inhibitor of ALK phosphorylation at this dose and time point (42) (**Fig. 1B**), which would not be affected by concomitant *RAS* pathway mutation (43).

To extend this work, we screened a selection of five neuroblastoma PDX-derived *ex-vivo* tumor cell models (PDTCs) carrying *ALK* alterations: GR-NB4 carried *ALK* amplification and IC-pPDX-112 carried an intron 3 *ALK* amplification (**Supplementary Fig. S1B**), and IC-pPDX-75, HSJD-NB-011 and HSJD-NB-012 carried *ALK*^F1174L^, *ALK*^I1171N^ and *ALK*^F1174C^ mutations, respectively (**Supplementary Table S2**). None of these models had concomitant RAS-MAPK pathway alteration. Notably, the patient from whom the IC-pPDX-75 model was derived had received crizotinib treatment for 12 months before the establishment of the PDX model.

PDTCs were treated with a panel of four ALK inhibitors (**Fig. 1Ci-iv, Supplementary Fig. S2B**). Drug responses were represented by the quantitative drug sensitivity score (DSS) (36), which integrates a multiparameter analysis, including the potency (the half-maximal effective concentration, EC_50_), slope of the dose–response curve, the area under the curve (AUC), and the maximum effect of the drug. The correlation between two experimental repeats (R1 and R2) was good across all four models (**Supplementary Fig. S2A**). In PDTC models, while DSS values of lorlatinib were higher than those of ceritinib or similar to those of crizotinib, only lorlatinib showed significantly higher cytotoxicity (lower DSS values) in *ALK*-aberrant PDTC models compared to wild-type models. Altogether our data supports previous work that lorlatinib is the most potent ALK inhibitor (24, 25). However, its efficacy in *in vitro* neuroblastoma models is not superior to other ALK inhibitors when a concomitant RAS/MAPK pathway mutation is present.

To evaluate the *in vivo* efficacy of ALK inhibitors, we performed a short-term response assessment study in the Th-*ALK*^F1174L^/*MYCN* GEMM model of *ALK* mutant neuroblastoma. Th-*ALK^F1174L^/MYCN* double transgenic animals develop spontaneous abdominal neuroblastoma at a shorter latency and 100% penetrance, compared to their Th-*MYCN* littermates, indicating that the addition of *ALK^F1174L^* plays a key role in tumorigenesis in this model (44). Tumor-bearing Th-*ALK*^F1174L^/*MYCN* mice, which had been treated with lorlatinib showed the greatest reduction in tumor volume compared with other clinical ALK inhibitors, over a 3-day interventional dosing schedule (**Fig. 2A**). When tumors were harvested for immunoassay measurement of total and pY1586 ALK, it was found that alectinib had an equivalent effect to lorlatinib on reduction of pY1586 / total ALK in the tumor cells, suggesting that the tumor volume response observed in these animals following lorlatinib treatment, may be due to a secondary off-target effect of lorlatinib, not shared with alectinib (**Fig. 2B**). Treatment of Th-*MYCN* heterozygous mice with crizotinib and alectinib also resulted in a highly variable tumor volume response over three days of therapy, confirming the activity of these compounds against targets other than mutant ALK (**Supplementary Fig. S3A**). Subsequently, survival of the Th-*ALK^F1174L^/MYCN* GEMM treated continuously with lorlatinib twice daily (BD), a schedule suggested by previous pharmacokinetic studies in mice (45), was assessed against vehicle control (**Fig. 2C**). Tumor volume measurements using MRI indicated that lorlatinib only slowed tumor progression in this model, with no tumor-free survivors observed (**Fig. 2D and E**). Analysis of tumor tissue taken at the end of the study revealed sustained ALK dephosphorylation and inhibition of expression of endogenous murine *mycn* as a downstream marker of ALK activity (44), suggesting that lorlatinib was still active against ALK signaling in these tumors (**Supplementary Fig. S3B and C**). Furthermore, RNA sequencing of tumor tissue taken at the end of the experiment, comparing vehicle treated controls against lorlatinib treated Th-*ALK^F1174L^/MYCN*, revealed reduced expression of noradrenergic genes, which was associated with lower ALK expression (**Supplementary Fig S3E**). The adrenergic phenotype has been associated with ALK expression in neuroblastoma, suggesting that lorlatinib treatment either allows expansion of non-ALK expressing cells or that ALK inhibition can drive a cell plasticity response (46). Additionally, there was an enhanced reduction of ALK expression in tumors classified as “poor responders” compared to “early responders”, based upon evidence of tumor regression at day 7 of the study (**Supplementary Fig. S3D and E**) (47).

**Figure 2.**
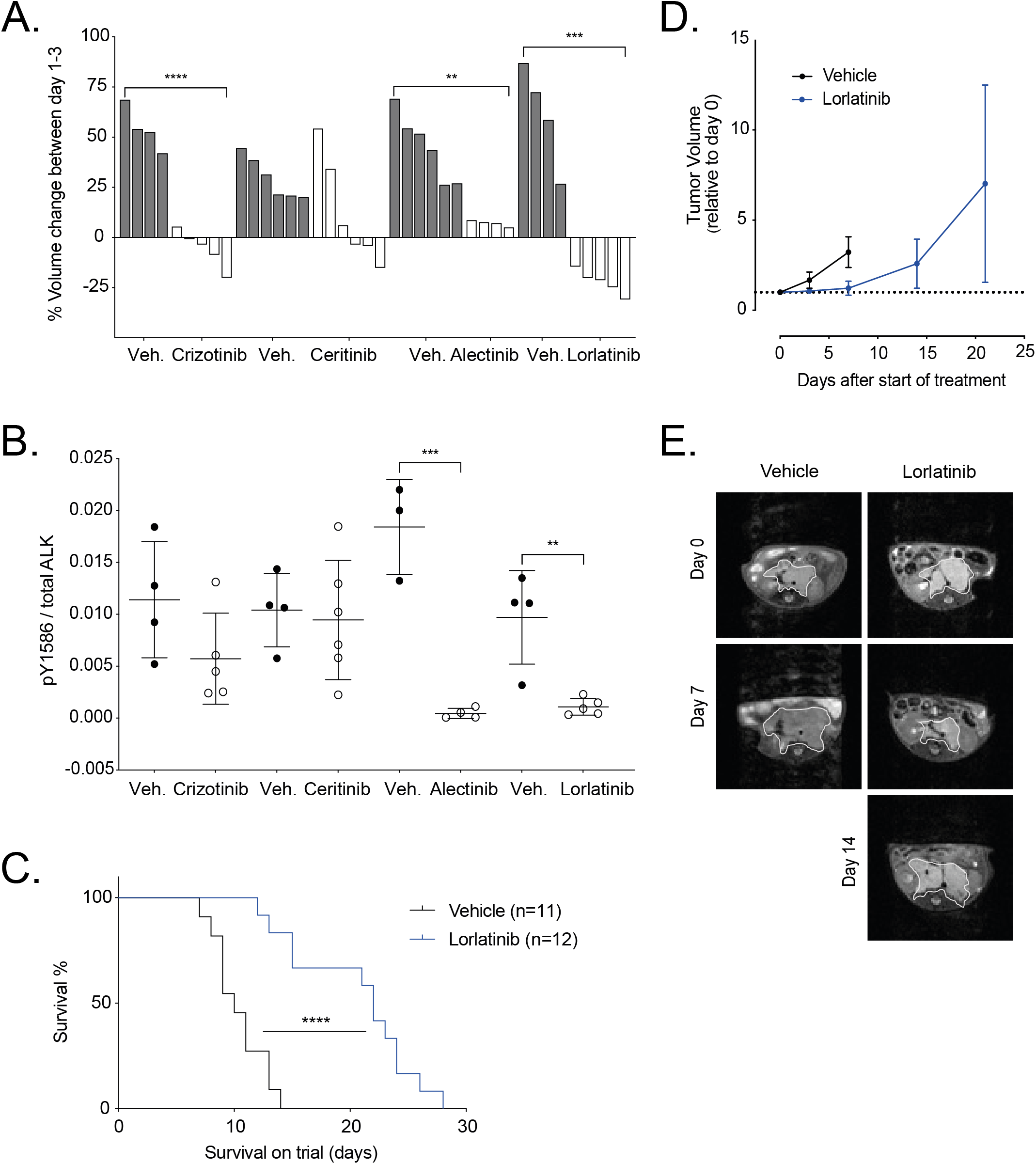
Lorlatinib treatment of Th-*ALK^F1174L^/MYCN* tumor-bearing animals gives a survival advantage over vehicle control. (A) *In vivo* analysis of a panel of ALK inhibitors including crizotinib, ceritinib, alectinib and lorlatinib was carried out using the Th-*ALK*^F1174L^/*MYCN* model. Tumor-bearing Th-*ALK*^F1174L^/*MYCN* mice, were treated with the indicated inhibitor or it’s corresponding vehicle (veh.) over a 3-day interventional dosing schedule, and tumor volume change was monitored by MRI on day 0 and day 3. Each bar represents tumor volume change in an individual animal. Crizotinib versus vehicle p<0.0001****. Alectinib versus vehicle p=0.0021**. Lorlatinib versus vehicle p=0.0002***. (B) Tumors from (A) were harvested for immunoassay testing of ALK and pY1586 ALK status. Alectinib versus vehicle p=0.0005***. Lorlatinib versus vehicle p=0.0037**. (C) Th-*ALK*^F1174L^/*MYCN* animals were treated with lorlatinib BD versus vehicle control to assess survival. P<0.0001**** according to Log-rank (Mantel-Cox) test. (D) Tumor volume was monitored by MRI. (E) Representative abdominal MRI of an animal from the lorlatinib survival study versus vehicle control. Tumor outlined by white line.

### Lorlatinib combined with chemotherapy improves tumor growth control in immunocompetent GEMM

We next evaluated whether concurrent ALK inhibition could increase the efficacy of conventional multi-agent chemotherapy regimens representative of those used in clinical practice. To this end we developed a multi-agent chemotherapy regime, consisting of vincristine, doxorubicin and cyclophosphamide (CAV), given as a single dose, prior to commencement of once daily (OD) lorlatinib (41). We employed this schedule in the TH-*ALK^F1174L^/MYCN* and TH-*MYCN* GEMM and found that the addition of lorlatinib to CAV chemotherapy significantly increased survival in this model, but not in the heterozygous Th-*MYCN* GEMM (**Fig. 3A**). Notably, animals treated with lorlatinib alone exhibited no survival advantage against vehicle controls with the once daily dosing schedule. Animals receiving the combination therapy underwent an early tumor response to treatment as measured by MRI (**Fig. 3B**), but ultimately no tumor-free survivors were observed.

**Figure 3:**
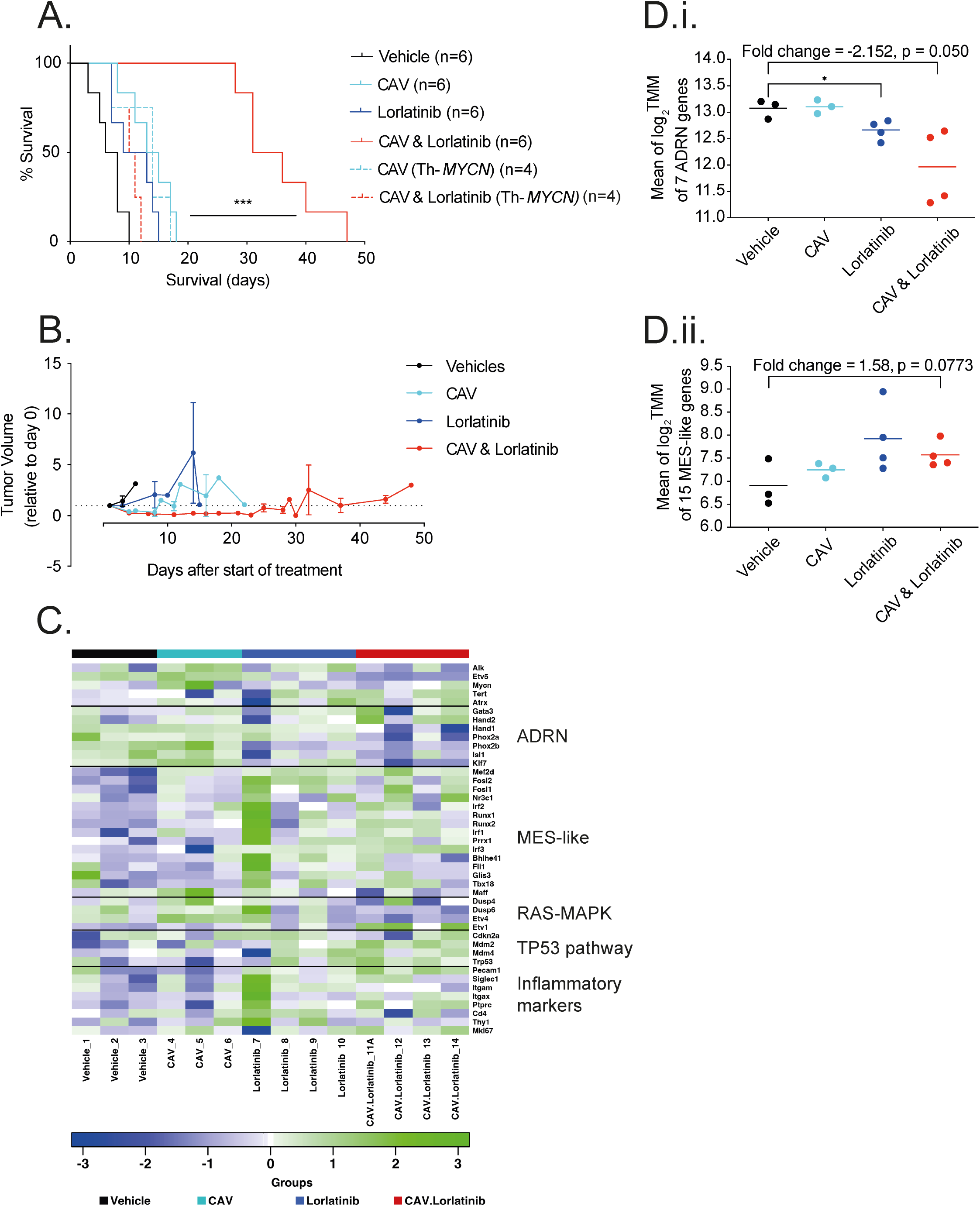
CAV chemotherapy in combination with lorlatinib leads to significantly enhanced survival in preclinical Th-*ALK*^F1174L^/*MYCN* GEMM neuroblastoma. (A) survival study of lorlatinib, with and without CAV chemotherapy in Th-*ALK*^F1174L^/*MYCN* and Th-*MYCN* (dashed line) GEMMs. Vehicle versus CAV and lorlatinib p=0.0006***; CAV versus CAV and lorlatinib p=0.0005***; lorlatinib versus CAV and lorlatinib p=0.0005*** in the Th-*ALK^F1174L^/MYCN* model according to log-rank (Mantel-Cox) test. (B) MRI growth monitoring of Th-*ALK^F1174L^/MYCN* tumors from the survival study in (A). No statistical significance between groups. (C) Heatmap of the expression of 34 selected genes across all the treatment groups. Noradrenergic (ADRN) and MES-like (mesenchymal cell - like). (D) Mean of log_2_ TMM (trimmed mean of M values) score for genes in ADRN (i) or MES (ii) panel across treatment groups.

We carried out RNA sequencing of tumors from the endpoint of this CAV-lorlatinib survival study to further investigate the reason for development of resistance on continuous daily dosing of lorlatinib. The greatest difference in overall gene expression was found between the vehicle-treated and combination-treated arms, within which we found upregulation of 135 genes and down regulation of 228 genes in tumors treated with CAV-lorlatinib (**Supplementary Fig S4A**). Furthermore we considered *ALK*-signature genes in this experimental cohort, and found that whilst there was no significant change in the mean *ALK* gene set expression, individually the expression of *ETV5*, known to be a robust marker of ALK activity, was significantly lowered between vehicle and combination-treated tumors (**Supplementary Fig. S4B.i and ii**) (48, 49). As in the earlier survival study comparing single agent lorlatinib against vehicle (**Supplementary Fig. S3E**) we found a trend towards expression of mesenchymal-like genes, rather than noradrenergic genes in both lorlatinib and combination-treated arms (**Fig. 3C and D; Supplementary Fig. 4C**), in keeping with recently published data (46). Additionally, in both the lorlatinib and lorlatinib-chemotherapy arms we found activation of the Tp53 pathway according to analysis of gene expression (**Fig. 3C**), consistent with previous studies showing activation of the TP53 pathway after ALK inhibition and chemotherapy (21). However, there was no apparent upregulation of RAS/MAPK associated genes in the combination treated group, activation of which is also associated with neuroblastoma relapse and resistance to ALK inhibition.

To expand our study of CAV in combination with lorlatinib *in vivo*, we used the identical dosing schedule as developed for Th-*ALK^F1174L^/MYCN* tumor-bearing animals in our neuroblastoma PDX panel. We employed this dosing schedule in an *ALK*-amplified and intron 3 *ALK*-amplified and one *ALK*-mutant (*ALK*^F1174C^) PDX models (**Supplementary Table S2**). Pharmacodynamic studies were performed by treating a cohort of PDX up until day 3. We analyzed levels of total ALK, P53, cleaved caspase 3 and P21 expression by immunohistochemistry in PDX samples in treated and vehicle-treated tumors at D3 (**Supplementary Table S3**). The GR-NB4 model exhibited the highest levels of ALK mRNA and protein expression in vehicle-treated tumors amongst the three models (**Fig. 4.A and Supplementary Fig. S5A**) However overall, the level of ALK protein expression was otherwise stable at this time point in all PDX models independently of the treatment received. No differences in cleaved caspase-3 and Ki67 protein expression were observed in *ALK*-amplified and intron 3 ALK-amplified PDX models in untreated and lorlatinib or lorlatinib-CAV treated PDX at this early time point (**Supplementary Table S3**). Of note GR-NB4 had very low *CDKN2A* expression (**Supplementary Fig. S5A**) consistent with *CDKN2A* homozygous deletion (**Supplementary Table S2**).

**Figure 4:**
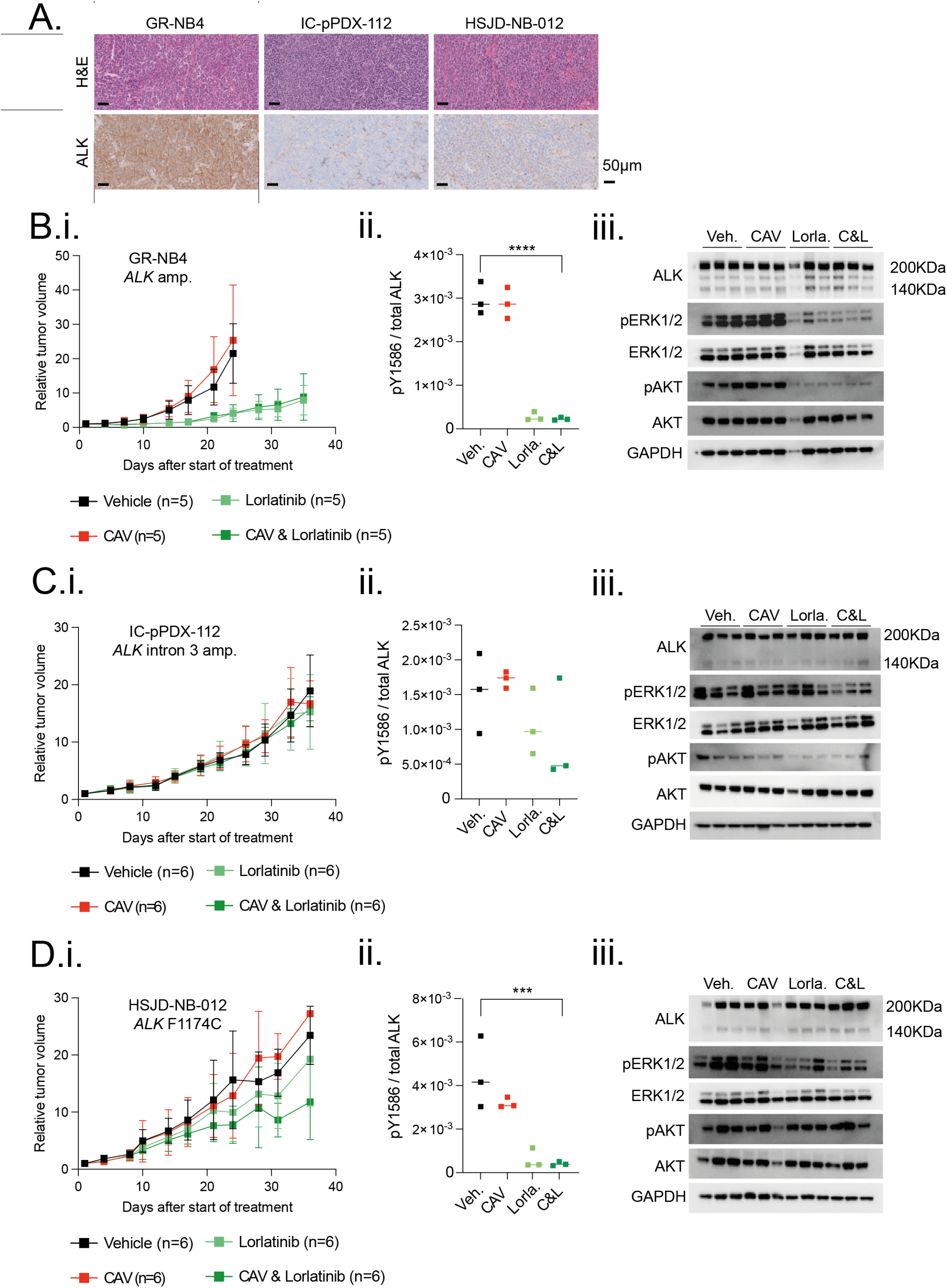
High baseline *in vivo* expression of ALK determines sensitivity to lorlatinib in an *ALK*-amplified PDX neuroblastoma model. (A) Hematoxylin and Eosin (H&E) and ALK immunohistochemical staining of vehicle-treated tumors from indicated PDX models, treated for 3 days. GR-NB4 (B), IC-pPDX-112 (C) and HSJD-NB-012 (D) PDX models were treated with vehicle control, one dose of chemotherapy (CAV: Vincristine, doxorubicin and cyclophosphamide), continuous lorlatinib, or chemotherapy and continuous lorlatinib combination. (i) Tumor volumes were monitored during treatment. (ii) pY1586/total ALK measured by immunoassay in tumor lysates taken at the end of the experiment (mean of 2 technical replicates) (Veh.: vehicle; Lorla.: Lorlatinib; C&L: CAV and lorlatinib). One-way ANOVA: B.ii: p<0.0001;C.ii: not significant; D.ii: p=0.0010 (iii) Immunoblots of signaling pathways downstream of ALK from tumor lysates as per (ii).

A longer-term tumor volume study was undertaken in a second cohort of mice. Total ALK expression was compared across the three models by immunoblot and immunoassay in vehicle treated tumors, confirming the highest level of ALK expression in the *ALK*-amplified GR-NB4 model (**Supplementary Fig. S5Bi and ii**). In this *ALK*-amplified GR-NB4 model, lorlatinib alone showed significant activity (TGI 99% at 24 days) (**Fig. 4B.i**). The addition of CAV did not significantly increase the efficacy of lorlatinib alone. Conversely, there was no effect on tumor growth in the intron 3 *ALK* amplified model (**Fig.4Ci**). The *ALK*-mutant PDX model showed a modest response to lorlatinib, both alone, and combined with CAV (**Fig. 4Di**). ALK immunoassay analysis of tumor lysates taken at the end of these experiments (**Fig. 4B-Dii**) revealed robust dephosphorylation of pY1586 ALK in GR-NB4 and HSJD-NB-012 but not IC-pPDX-112. There was good evidence of dephosphorylation of both AKT and ERK1/2 on immunoblotting following lorlatinib treatment in the GR-NB4 model, and dephosphorylation limited to ERK1/2 in the IC-pPDX-112 model (**Fig. 4B-Diii, Supplementary Fig. S5Ci-iii**). Furthermore, studying the *ALK* expression intensity evaluated by RNA sequencing in 44 patients with neuroblastoma from the MAPPYACTS trial (50), indicated that mutation or amplification of ALK is associated with an increased RNA expression of ALK, in comparison to non-ALK aberrant neuroblastoma (**Supplementary Fig. 5D**).

### Idasanutlin synergizes with ALK inhibitors in TP53 wild-type and ALK aberrant neuroblastoma cell lines

Whilst the combination of lorlatinib with chemotherapy will be soon be introduced into upfront therapy for high-risk *ALK*-aberrant neuroblastoma, other combinations that might be suitable for this sub-group of patients following relapse and chemotherapy-resistance are still warranted. To this end, we also investigated combinations of ALK inhibitors with MDM2 inhibitors, which have previously been suggested as potential strategies to overcome resistance to ALK inhibitors (28, 29). Since none of the PDXs had RAS-MAPK mutations, this was not a mechanism of resistance to ALK inhibitors in these models and all of them were TP53 wild-type (**Supplementary Table S2**). The MDM2 antagonist, idasanutlin, is being evaluated in combination with venetoclax or chemotherapy for patients with relapsed/refractory neuroblastoma in a Phase I clinical trial (NCT04029688). In TP53-wild-type and ALK-aberrant neuroblastoma cell lines including a previously unreported *MDM2* and *ALK* amplified cell line, NB1691 with MDM2 amplification (**Supplementary Fig. S1A**) 2 *ALK* mutant and 2 *ALK* mutant and amplified cell lines (**Supplementary Table S1**). Median-effect-analysis demonstrated a synergistic interaction between idasanutlin and ALK inhibitors, TAE-684 and alectinib *in vitro* (**Fig. 5A & B**). Further assessment showed that the combination treatment led to enhanced levels of apoptosis as evident by the greater proportion of sub-G1 events and caspase 3/7 activity determined using FACS and caspase 3/7 assays (**Fig. 5C-F, Supplementary Fig. S6**).

**Figure 5:**
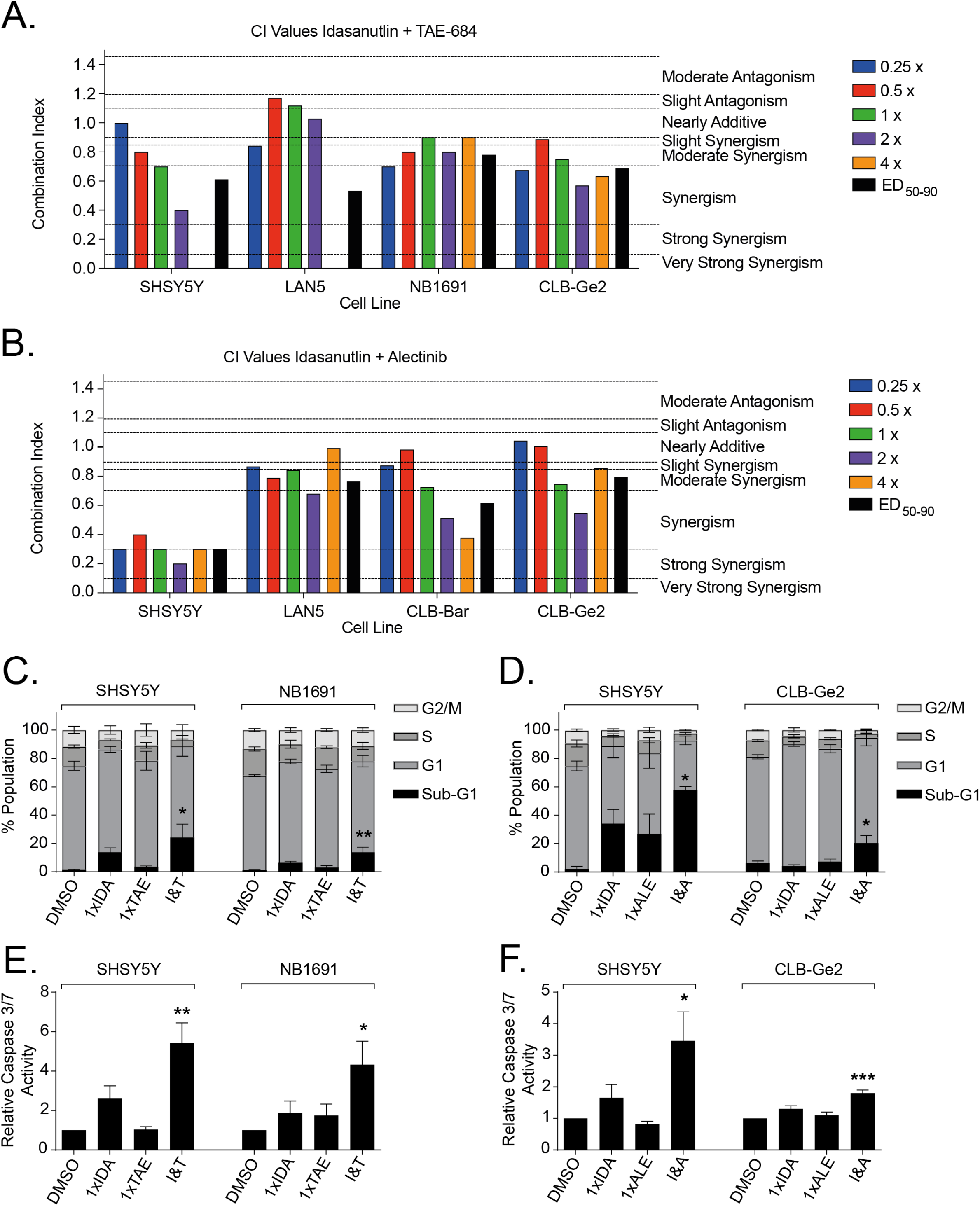
Idasanutlin synergizes with ALK inhibitors in *TP53* wild-type and *ALK* aberrant neuroblastoma cell lines. CI values at each constant 1:1 ratio combination and average of CI values at ED50, ED75 and ED90 of Idasanutlin in combination with A) TAE-684 and B) Alectinib. Functional analysis of Idasanutlin in combination with TAE-684 and Alectinib using sub-G1 and cell cycle phase distribution (C & D), and caspase 3/7 activity (E & F). IDA, Idasanutlin; TAE, TAE-684; I&T, Idasanutlin and TAE-684; ALE, Alectinib; I&A, Idasanutlin and Alectinib. One-way ANOVA: (C): SHSY5Y *p=0.0441; NB1691**p=0.0070; (D): SHSY5Y *p=0.0273; CLB-Ge2 *p=0.0230; (E): SHSY5Y **p=0.0043; NB1691 *p=0.0322; (F): SHSY5Y *p=0.0150; CLB-Ge2 ***p=0.0008.

### Idasanutlin enhances antitumor activity of lorlatinib in an ALK-amplified neuroblastoma PDX model

*In vivo* we studied the efficacy of lorlatinib in combination with idasanutlin in two *ALK*-amplified and one *ALK*-mutant PDX model. In the high-ALK expressing *ALK*-amplified GR-NB4 model, whereas idasanutlin alone had a modest effect on TGI (38% at 24 days), the combination of lorlatinib and idasanutlin showed impressive additive effect with complete remission (TGI 50% and 0% at 24 days for combination and inhibitors alone respectively) (**Fig. 6Ai-ii, Supplementary Table S4**). The animals in this combination arm were treated until day 84 and then left until tumor relapse occurred and the tumors reached the ethical size limit (**Supplementary Fig. S6**). Conversely, the *ALK* mutant model was completely resistant to both drugs as monotherapy as well as the combination (**Fig. 6Ci**). Immunoassay analysis for pY1586ALK expression from the samples at the end of the experiment on day 35-40 revealed robust dephosphorylation of ALK in most tumors receiving either lorlatinib alone or in combination with idasanutlin (**Fig. 6A-Cii**). The exception was from the lorlatinib-idasanutlin arm of the GR-NB4 model, as all animals had survived several weeks following termination of treatment (**Fig. 6Aii, Supplementary Fig. 6**). Pharmacodynamics were performed on day 3 on a separate cohort of animals. No differences in cleaved caspase-3 and Ki67 protein expression were observed in *ALK*-amplified and *ALK* mutant PDX models in samples treated with lorlatinib alone and in combination with idasanutlin (**Supplementary Table S3**). The level of ALK, TP53 and P21 expression by IHC in *ALK*-amplified model was unchanged in treated and untreated tumors.

**Figure 6:**
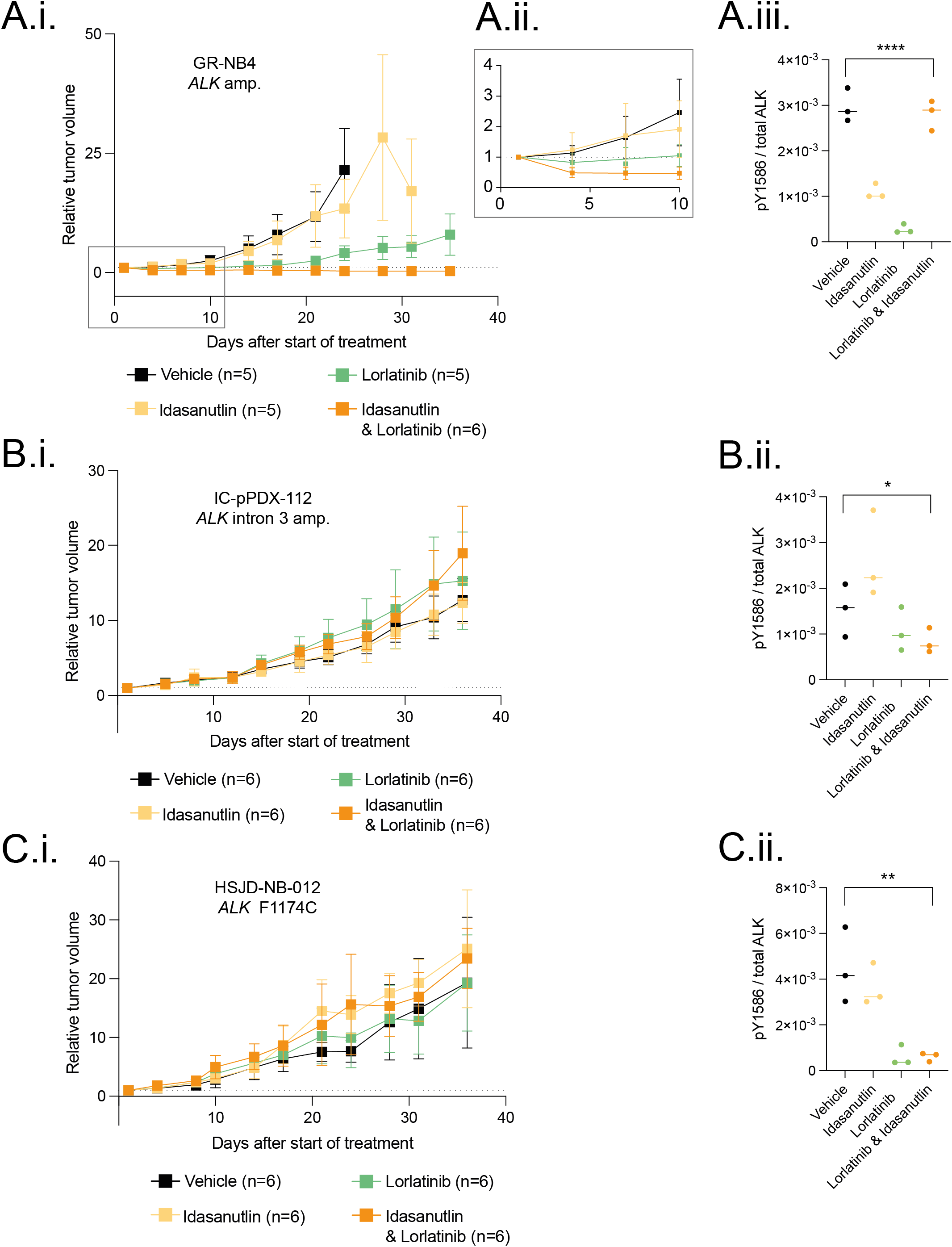
Combination effects of lorlatinib and idasanutlin in *ALK*-altered neuroblastoma PDX models. (A) GR-NB4 (B) IC-pPDX-112 (C) HSJD-NB-012 PDX models were treated with vehicle control, idasanutlin, lorlatinib, or idasanutlin and lorlatinib combination. (i) Tumor volumes were monitored during treatment (with (A.ii) inset of GR-NB4 PDX up to day 10). (A.iii, B.ii and C.ii) pY1586/total ALK measured by immunoassay in tumor lysates taken at the end of the experiment (mean of 2 technical replicates) One-way ANOVA: A.iii: p=<0.0001; B.ii: p=0.0331; C.ii: p=0.0020.

Bulk RNA-sequencing of representative PDX tumors from the experiments shown in Fig. 4A-C and Fig. 6A-C was undertaken, illustrating differences in gene expression across the 3 models (**Supplementary Fig. S5A**). This showed increased TP53 and MDM2 expression following combination treatment in IC-pPDX-112 consistent with functional TP53, but not in the GR-NB4 or HSJD-NB-012 PDX models.

## DISCUSSION

The goal of this study was to provide robust preclinical insights into the activity of ALK inhibitors in combination with chemotherapy or targeted MDM2 inhibition in neuroblastomas carrying mutations or amplifications of the *ALK* gene.

The ongoing International Society of Paediatric Oncology European Neuroblastoma (SIOPEN) trial HRNBL2 (NCT04221035) for first-line treatment of high-risk neuroblastoma will soon be amended to introduce lorlatinib in upfront treatment of patients with *ALK*-aberrant high-risk neuroblastoma. Concurrently, the Children’s Oncology Group is amending the ANBL1531 phase III trial to replace crizotinib with lorlatinib. Since 2008, when the role of *ALK* in neuroblastoma was first described (12, 13, 51, 52), the study of ALK inhibitors in neuroblastoma has grown as a result of progress in the development of these drugs for adult malignancies and advances made by the scientific neuroblastoma community.

Preclinical studies in *ALK*-aberrant neuroblastomas have shown intrinsic resistance of *ALK*^F1174^ and *ALK*^F1245^, two of the 3 main mutational hotspots, to crizotinib (18). The second-generation molecule ceritinib inhibits the kinases of ALK F1245 and ALK F1275 but also exhibits intrinsic resistance to the F1174 variant (23). Lorlatinib, a third generation ALK inhibitor overcomes resistance to first and second generation ALK inhibitors by its macrocyclic structure with optimized physicochemical properties, which are associated with improved metabolic stability, blood-brain barrier permeability and low propensity for multi-drug resistance (MDR) efflux (53).

Cell lines with *ALK* alterations were in general less sensitive to lorlatinib than to ceritinib and other ALK inhibitors based on growth inhibition assays (**Fig. 1Ai-v**) which we believe is due to the presence of a concomitant RAS-MAPK pathway mutation in cell lines which were hyper-resistant to lorlatinib (except for cell line NB1691, which has a highly complex genotype (**Supplementary Fig. S1A**)). Despite these GI_50_ data in cell lines, albeit with an assay measuring metabolic activity as a surrogate for cell viability, lorlatinib showed the greatest inhibition of ALK phosphorylation in cell lines, (**Fig. 1B**) which reflects its selectivity for the ALK signaling pathway upstream of the RAS-MAPK pathway compared with other ALK inhibitors, but inhibition of ALK signaling by lorlatinib can be by-passed in the presence of a downstream RAS-MAPK pathway mutation. Indeed, lorlatinib showed the greatest *in vivo* reduction in tumor volume over a 3-day interventional dosing schedule in *ALK*^F1174L^/*MYCN* GEMMs which do not have concomitant RAS-MAPK pathway mutations. However, alectinib had a similar effect to lorlatinib on ALK dephosphorylation in tumor cells, suggesting other off-target effects of lorlatinib which could lead to enhanced efficacy compared to alectinib (**Fig. 2A and B**). Furthermore, whereas CAV chemotherapy or lorlatinib alone did not show any survival advantage, the association of both was synergistic and induced significantly increased survival in the Th-*ALK*^F1174L^/*MYCN* neuroblastoma model against continuous once daily lorlatinib or one dose of CAV alone (**Fig. 3A**). Additionally, our observation that GEMM tumors relapsing during treatment with lorlatinib alone (**Fig. 2C**) or the CAV lorlatinib combination (**Fig. 3A**) exhibit a shift from the adrenergic state, is in keeping with recently published data describing ALK inhibitor resistance of mesenchymal-type neuroblastoma xenografts which express low levels of ALK (46). We can therefore speculate that either there was a subpopulation of tumor cells inherently resistant to therapy or that phenotypic plasticity might have played a role in the acquisition of resistance to ALK inhibition and chemotherapy in our GEMM model.

Interest in PDXs has increased in recent years due to the development of personalized cancer medicines based on genomic profiling and they have improved the predictive power of preclinical therapeutic studies (43). From *ex vivo* screening of PDTC models, whereas ceritinib was more active and crizotinib showed similar activity to lorlatinib in ALK-aberrant models, lorlatinib was the only ALK inhibitor to show statistically significant enhanced cytotoxicity in ALK-aberrant models compared to wild-type, reinforcing the hypothesis of off-target effects of ceritinib or crizotinib (**Fig. 1Ci-iv**). *In vivo* studies showed high efficacy of lorlatinib alone in the GR-NB4 *ALK*-amplified model, where the addition of chemotherapy did not increase the tumor growth inhibition and chemotherapy alone was ineffective (**Fig. 4Ai**). This may be due to the low doses of chemotherapy chosen for these experiments, as CAV doses were low to help reveal a potential synergy between chemotherapy and lorlatinib. It is also possible that an immunocompetent mouse model might be required to demonstrate preclinical efficacy of chemotherapy via immunostimulation, therefore the study of the microenvironment in our GEMM is ongoing. Unexpectedly, the *ALK*^F1174C^ mutated model (HSJD-NB-012) exhibited only a modest response to lorlatinib alone and in combination with chemotherapy (**Fig. 4Di**), the reasons for which are currently unclear. The lack of response by the PDX with a “silent” Intron 3 ALK amplification (IC-pPDX-112), as opposed to amplification across the entire coding sequence, fits with the relatively low expression of ALK RNA and protein. The high mutational burden in this model, which is unusual for a diagnostic neuroblastoma, and could give rise to especially aggressive tumor growth.

In view of the future amended HRNBL2 protocol combining upfront lorlatinib with chemotherapy upfront in *ALK*-aberrant tumors from high-risk neuroblastoma patients, we investigated the possibility of combined ALK and MDM2 inhibition for *ALK*-aberrant relapsed neuroblastoma, as ALK and MDM2 inhibitors have already been explored as potential strategies to overcome resistance to ALK inhibitors (28, 29), and TP53 mutations are rare in relapsed neuroblastoma. In several *TP53*-wild-type and *ALK*-aberrant (mutated, amplified or both) cell lines, the combination of idasanutlin and ALK inhibitors was found to be synergistic and induced enhanced levels of apoptosis (**Fig. 5**). In the *ALK*-amplified GR-NB4 PDX model, the combination of lorlatinib and idasanutlin showed impressive additive effect with complete remission (**Fig. 6Ai**), which may be partly explained by the additional presence of a homozygous CDKN2A/B deletion in this model. Although it was not possible to test idasanutlin, which is specific for human MDM2 (54) in the Th-*ALK^F1174L^/MYCN* GEMM, our previous studies have shown that *ALK^F1174L^* promotes anti-apoptotic activity in this model of neuroblastoma (44). Therefore, future work to investigate whether a mouse reactive MDM2 inhibitor is synergistic with ALK inhibition via enhanced apoptosis is warranted.

Importantly our comprehensive study of ALK inhibitor combinations included a pharmacodynamic assessment of ALK expression, ALK phosphorylation and down-stream activity of ALK across our models. We found a high baseline level of tumoral ALK expression was associated with sensitivity to ALK inhibition. This finding supports measurement of total ALK expression alongside *ALK* aberrations in future clinical studies with ALK inhibitors to determine if there is a threshold of ALK expression that might predict a response to ALK inhibitor combinations in *ALK* aberrant neuroblastoma.

## Acknowledgements

The authors would like to thank Pfizer for providing lorlatinib for the *in vivo* GEMM work, and Roche-Genentech for providing both idasanutlin for the *in vitro* work and alectinib for the *in vivo* GEMM work. We wish to thank the Biological Services Unit at The Institute of Cancer Research (London), and the Animal Platform of the Institut Curie-Orsay (Paris), as well as Adnan Naguez and Marine Michou for their technical assistance.

## Data Availability Statement

The data generated in this study are publicly available at BioProject, ID PRJNA848127.

## Notes

**ADDITIONAL INFORMATION**, **Financial support:** In the UK, ERT, EP, HC, BMC, KB, SS and LC were supported by a Cancer Research UK (CRUK) Programme Grant (A28278); EC was supported by a Children with Cancer Clinical PhD Fellowship (CWL022X); QG was supported by a Cancer Research UK (A24566); YJ was supported by a Children with Cancer UK Research Fellowship (2014/176). SG was supported by Royal Marsden NIHR Biomedical Research Centre funding, a CRUK Postdoctoral Research Bursary and a CRUK Clinical Scientist Fellowship. DT, LC, SD, and AG received funding from Neuroblastoma UK, Niamh’s Next Step, Children’s Cancer North, the Newcastle National Institute for Health Research Biomedical Research Centre and the Children’s Cancer & Leukaemia Group/Little Princess Trust. In France, this work was supported by the Annenberg Foundation, the Association Hubert Gouin Enfance et Cancer, the Fédération Enfants Cancers Santé, the Société Française de lutte contre les Cancers et les leucémies de l’Enfant et l’adolescent (SFCE), Les Bagouz à Manon, Les amis de Claire and the Fondation ARC pour la Recherche contre le Cancer (ARC). Funding was also obtained from SiRIC/INCa (Grant INCa-DGOS-4654), and PHRC IC2007-09 grant. High-throughput sequencing was performed by the ICGex NGS platform of the Institut Curie supported by the grants ANR-10-EQPX-03 (Equipex) and ANR-10-INBS-09-08 (France Génomique Consortium) from the Agence Nationale de la Recherche (“Investissements d’Avenir” program), by the Canceropole Ile-de-France and by the SiRIC-Curie program - SiRIC Grant “INCa-DGOS-4654“. Tumor sequencing data and clinical data have been provided under the MAPPYACTS protocol (clinicaltrial.gov: NCT02613962). The drug screening was supported in the framework of ERA PerMed (ERAPERMED2018-121, COMPASS). Molecular characterization of the PDX models was supported within the framework of the IMI2 ITCC P4 program funded by IMI2 grant agreement no. 116064 ITCC-P4. BG is supported by the Parrainage médecin-chercheur of Gustave Roussy.

**Conflict of interest disclosure statement:** The authors declare no potential conflicts of interest.

### Competing Interest Statement

The authors have declared no competing interest.

## REFERENCES

1. Brodeur GM. Spontaneous regression of neuroblastoma. Cell Tissue Res. 2018;372(2):277–86.

2. Pinto NR, Applebaum MA, Volchenboum SL, Matthay KK, London WB, Ambros PF, et al. Advances in Risk Classification and Treatment Strategies for Neuroblastoma. J Clin Oncol. 2015;33(27):3008–17.

3. Matthay KK, Maris JM, Schleiermacher G, Nakagawara A, Mackall CL, Diller L, et al. Neuroblastoma. Nat Rev Dis Primers. 2016;2:16078.

4. Brodeur GM, Seeger RC, Schwab M, Varmus HE, Bishop JM. Amplification of N-myc in untreated human neuroblastomas correlates with advanced disease stage. Science. 1984;224(4653):1121–4.

5. Ackermann S, Cartolano M, Hero B, Welte A, Kahlert Y, Roderwieser A, et al. A mechanistic classification of clinical phenotypes in neuroblastoma. Science. 2018;362(6419):1165–70.

6. Peifer M, Hertwig F, Roels F, Dreidax D, Gartlgruber M, Menon R, et al. Telomerase activation by genomic rearrangements in high-risk neuroblastoma. Nature. 2015;526(7575):700–4.

7. Valentijn LJ, Koster J, Zwijnenburg DA, Hasselt NE, van Sluis P, Volckmann R, et al. TERT rearrangements are frequent in neuroblastoma and identify aggressive tumors. Nat Genet. 2015;47(12):1411–4.

8. Cheung NK, Zhang J, Lu C, Parker M, Bahrami A, Tickoo SK, et al. Association of age at diagnosis and genetic mutations in patients with neuroblastoma. JAMA. 2012;307(10):1062–71.

9. Hartlieb SA, Sieverling L, Nadler-Holly M, Ziehm M, Toprak UH, Herrmann C, et al. Alternative lengthening of telomeres in childhood neuroblastoma from genome to proteome. Nat Commun. 2021;12(1):1269.

10. Bellini A, Potschger U, Bernard V, Lapouble E, Baulande S, Ambros PF, et al. Frequency and Prognostic Impact of ALK Amplifications and Mutations in the European Neuroblastoma Study Group (SIOPEN) High-Risk Neuroblastoma Trial (HR-NBL1). J Clin Oncol. 2021;39(30):3377–90.

11. Zeineldin M, Federico S, Chen X, Fan Y, Xu B, Stewart E, et al. MYCN amplification and ATRX mutations are incompatible in neuroblastoma. Nat Commun. 2020;11(1):913.

12. Janoueix-Lerosey I, Lequin D, Brugieres L, Ribeiro A, de Pontual L, Combaret V, et al. Somatic and germline activating mutations of the ALK kinase receptor in neuroblastoma. Nature. 2008;455(7215):967–70.

13. Mosse YP, Laudenslager M, Longo L, Cole KA, Wood A, Attiyeh EF, et al. Identification of ALK as a major familial neuroblastoma predisposition gene. Nature. 2008;455(7215):930–5.

14. De Brouwer S, De Preter K, Kumps C, Zabrocki P, Porcu M, Westerhout EM, et al. Meta-analysis of neuroblastomas reveals a skewed ALK mutation spectrum in tumors with MYCN amplification. Clin Cancer Res. 2010;16(17):4353–62.

15. Eleveld TF, Oldridge DA, Bernard V, Koster J, Colmet Daage L, Diskin SJ, et al. Relapsed neuroblastomas show frequent RAS-MAPK pathway mutations. Nat Genet. 2015;47(8):864–71.

16. Padovan-Merhar OM, Raman P, Ostrovnaya I, Kalletla K, Rubnitz KR, Sanford EM, et al. Enrichment of Targetable Mutations in the Relapsed Neuroblastoma Genome. PLoS Genet. 2016;12(12):e1006501.

17. Schleiermacher G, Javanmardi N, Bernard V, Leroy Q, Cappo J, Rio Frio T, et al. Emergence of new ALK mutations at relapse of neuroblastoma. J Clin Oncol. 2014;32(25):2727–34.

18. Bresler SC, Wood AC, Haglund EA, Courtright J, Belcastro LT, Plegaria JS, et al. Differential inhibitor sensitivity of anaplastic lymphoma kinase variants found in neuroblastoma. Sci Transl Med. 2011;3(108):108ra14.

19. Bourdeaut F, Ferrand S, Brugieres L, Hilbert M, Ribeiro A, Lacroix L, et al. ALK germline mutations in patients with neuroblastoma: a rare and weakly penetrant syndrome. Eur J Hum Genet. 2012;20(3):291–7.

20. Foster JH, Voss SD, Hall DC, Minard CG, Balis FM, Wilner K, et al. Activity of Crizotinib in Patients with ALK-Aberrant Relapsed/Refractory Neuroblastoma: A Children’s Oncology Group Study (ADVL0912). Clin Cancer Res. 2021;27(13):3543–8.

21. Krytska K, Ryles HT, Sano R, Raman P, Infarinato NR, Hansel TD, et al. Crizotinib Synergizes with Chemotherapy in Preclinical Models of Neuroblastoma. Clin Cancer Res. 2016;22(4):948–60.

22. Greengard E, Mosse YP, Liu X, Minard CG, Reid JM, Voss S, et al. Safety, tolerability and pharmacokinetics of crizotinib in combination with cytotoxic chemotherapy for pediatric patients with refractory solid tumors or anaplastic large cell lymphoma (ALCL): a Children’s Oncology Group phase 1 consortium study (ADVL1212). Cancer Chemother Pharmacol. 2020;86(6):829–40.

23. Fischer M, Moreno L, Ziegler DS, Marshall LV, Zwaan CM, Irwin MS, et al. Ceritinib in paediatric patients with anaplastic lymphoma kinase-positive malignancies: an open-label, multicentre, phase 1, dose-escalation and dose-expansion study. Lancet Oncol. 2021;22(12):1764–76.

24. Infarinato NR, Park JH, Krytska K, Ryles HT, Sano R, Szigety KM, et al. The ALK/ROS1 Inhibitor PF-06463922 Overcomes Primary Resistance to Crizotinib in ALK-Driven Neuroblastoma. Cancer Discov. 2016;6(1):96–107.

25. Guan J, Tucker ER, Wan H, Chand D, Danielson LS, Ruuth K, et al. The ALK inhibitor PF-06463922 is effective as a single agent in neuroblastoma driven by expression of ALK and MYCN. Dis Model Mech. 2016;9(9):941–52.

26. Debruyne DN, Dries R, Sengupta S, Seruggia D, Gao Y, Sharma B, et al. BORIS promotes chromatin regulatory interactions in treatment-resistant cancer cells. Nature. 2019;572(7771):676–80.

27. Berlak M, Tucker E, Dorel M, Winkler A, McGearey A, Rodriguez-Fos E, et al. Mutations in ALK signaling pathways conferring resistance to ALK inhibitor treatment lead to collateral vulnerabilities in neuroblastoma cells. Mol Cancer. 2022;21(1):126.

28. Miyazaki M, Otomo R, Matsushima-Hibiya Y, Suzuki H, Nakajima A, Abe N, et al. The p53 activator overcomes resistance to ALK inhibitors by regulating p53-target selectivity in ALK-driven neuroblastomas. Cell Death Discov. 2018;4:56.

29. Wang HQ, Halilovic E, Li X, Liang J, Cao Y, Rakiec DP, et al. Combined ALK and MDM2 inhibition increases antitumor activity and overcomes resistance in human ALK mutant neuroblastoma cell lines and xenograft models. Elife. 2017;6.

30. Carr-Wilkinson J, O’Toole K, Wood KM, Challen CC, Baker AG, Board JR, et al. High Frequency of p53/MDM2/p14ARF Pathway Abnormalities in Relapsed Neuroblastoma. Clin Cancer Res. 2010;16(4):1108–18.

31. Chen Z, Lin Y, Barbieri E, Burlingame S, Hicks J, Ludwig A, et al. Mdm2 deficiency suppresses MYCN-Driven neuroblastoma tumorigenesis in vivo. Neoplasia. 2009;11(8):753–62.

32. Swarbrick A, Woods SL, Shaw A, Balakrishnan A, Phua Y, Nguyen A, et al. miR-380-5p represses p53 to control cellular survival and is associated with poor outcome in MYCN-amplified neuroblastoma. Nat Med. 2010;16(10):1134–40.

33. Veschi V, Thiele CJ. High-SETD8 inactivates p53 in neuroblastoma. Oncoscience. 2017;4(3-4):21–2.

34. Chen L, Humphreys A, Turnbull L, Bellini A, Schleiermacher G, Salwen H, et al. Identification of different ALK mutations in a pair of neuroblastoma cell lines established at diagnosis and relapse. Oncotarget. 2016;7(52):87301–11.

35. Chen L, Rousseau RF, Middleton SA, Nichols GL, Newell DR, Lunec J, et al. Pre-clinical evaluation of the MDM2-p53 antagonist RG7388 alone and in combination with chemotherapy in neuroblastoma. Oncotarget. 2015;6(12):10207–21.

36. Stewart E, Federico SM, Chen X, Shelat AA, Bradley C, Gordon B, et al. Orthotopic patient-derived xenografts of paediatric solid tumours. Nature. 2017;549(7670):96–100.

41. Garaventa A, Poetschger U, Valteau-Couanet D, Luksch R, Castel V, Elliott M, et al. Randomized Trial of Two Induction Therapy Regimens for High-Risk Neuroblastoma: HR-NBL1.5 International Society of Pediatric Oncology European Neuroblastoma Group Study. J Clin Oncol. 2021;39(23):2552–63.

43. Tucker ER, George S, Angelini P, Bruna A, Chesler L. The Promise of Patient-Derived Preclinical Models to Accelerate the Implementation of Personalised Medicine for Children with Neuroblastoma. J Pers Med. 2021;11(4).

44. Berry T, Luther W, Bhatnagar N, Jamin Y, Poon E, Sanda T, et al. The ALK(F1174L) mutation potentiates the oncogenic activity of MYCN in neuroblastoma. Cancer Cell. 2012;22(1):117–30.

45. Yamazaki S, Lam JL, Zou HY, Wang H, Smeal T, Vicini P. Translational pharmacokinetic-pharmacodynamic modeling for an orally available novel inhibitor of anaplastic lymphoma kinase and c-Ros oncogene 1. J Pharmacol Exp Ther. 2014;351(1):67–76.

46. Westerhout EM, Hamdi M, Stroeken P, Nowakowska NE, Lakeman A, van Arkel J, et al. Mesenchymal-Type Neuroblastoma Cells Escape ALK Inhibitors. Cancer Res. 2022;82(3):484–96.

47. Boeva V, Louis-Brennetot C, Peltier A, Durand S, Pierre-Eugene C, Raynal V, et al. Heterogeneity of neuroblastoma cell identity defined by transcriptional circuitries. Nat Genet. 2017;49(9):1408–13.

48. Lambertz I, Kumps C, Claeys S, Lindner S, Beckers A, Janssens E, et al. Upregulation of MAPK Negative Feedback Regulators and RET in Mutant ALK Neuroblastoma: Implications for Targeted Treatment. Clin Cancer Res. 2015;21(14):3327–39.

49. Mus LM, Lambertz I, Claeys S, Kumps C, Van Loocke W, Van Neste C, et al. The ETS transcription factor ETV5 is a target of activated ALK in neuroblastoma contributing to increased tumour aggressiveness. Sci Rep. 2020;10(1):218.

50. Berlanga P, Pierron G, Lacroix L, Chicard M, Adam de Beaumais T, Marchais A, et al. The European MAPPYACTS Trial: Precision Medicine Program in Pediatric and Adolescent Patients with Recurrent Malignancies. Cancer Discov. 2022;12(5):1266–81.

51. George RE, Sanda T, Hanna M, Frohling S, Luther W, 2nd, Zhang J, et al. Activating mutations in ALK provide a therapeutic target in neuroblastoma. Nature. 2008;455(7215):975–8.

52. Chen Y, Takita J, Choi YL, Kato M, Ohira M, Sanada M, et al. Oncogenic mutations of ALK kinase in neuroblastoma. Nature. 2008;455(7215):971–4.

53. Zou HY, Friboulet L, Kodack DP, Engstrom LD, Li Q, West M, et al. PF-06463922, an ALK/ROS1 Inhibitor, Overcomes Resistance to First and Second Generation ALK Inhibitors in Preclinical Models. Cancer Cell. 2015;28(1):70–81.

54. Chen L, Esfandiari A, Reaves W, Vu A, Hogarty MD, Lunec J, et al. Characterisation of the p53 pathway in cell lines established from TH-MYCN transgenic mouse tumours. Int J Oncol. 2018;52(3):967–77.

