## Supplementary data for "COMBINATION THERAPIES TARGETING ALK-ABERRANT NEUROBLASTOMA IN PRECLINICAL MODELS"

**SUPPLEMENTARY METHODS**

**Human Neuroblastoma cell lines**

The sources of the cell lines were as follows: CLB-Ge2, CLB-BAR-Rec (Valérie Combaret), IMR32 (Penny Lovat), SHSY5Y, SKNBE2c (Barbara Spengler), LAN5, LAN6, CHLA136 (Pat Reynolds), NB69 (John Maris), NBLW, NBLW-R (Sue Cohn), GIMEN (Mirco Ponzoni), LAN1, SKNAS (Jean Bénard), NB-1 (Katleen De Preter), NB1691 (Clinton Stewart), NGP (Rogier Versteeg), TR14 (Maria Łastowska).

**Determination of PDX genomic alterations**

*Sequencing. ALK*-aberrant PDX models are described in Supplementary Table 2. All PDX tumors and paired patient germline DNA were whole-exome sequenced (WES) by Agilent SureSelect Human All Exon v5 or Nimblegen Roche Sequencing SeqCap EZ MedExome Kit according to manufacturer’s protocols (paired-ends 100x100 bp, expected coverage 100 X).

*Bioinformatics sequencing pipeline.* For PDX samples, fastq files were aligned using Bwa-mem on a hybrid GRCh37-GRCm38 (human-mouse) reference genome. Reads exclusively mapped to human chromosomes with mapping quality more than 20 were extracted. For human samples, fastq files were aligned on GRCh37 human genome reference and uniquely mapped reads with mapping quality more than 20 were extracted. Finally, duplicate reads were removed. Coverage analysis was done using MOSDEPTH and GATK4. For copy number analysis, Sequenza and FACETS were used, and purity and the ploidy of each sample was extracted from Sequenza results. Mutect2 was used for variant calling. The mutations were filtered using Filter Mutect2 calls and annotated using VEP. Only the variants matching the following criteria were retained: the coverage should be more than 20X, variant supporting reads should be more than 10 in tumor and cfDNA with at least one forward and one reverse read, less than 2 reads in germline sample, VAF >0.01, predicted as deleterious by SIFT or PolyPhen, and frequency less than 0.001 in the population.

#### **mRNA Sequencing**

Excised tumors from GEMM and PDX mice that were enrolled on 4 different treatment arms: Vehicle, lorlatinib/Vehicle, CAV/Vehicle, CAV and lorlatinib combination therapy (CAV/Lorlatinib), were used for a gene expression study. Libraries were created using DNBSEQ Eukaryotic Strand-specific mRNA library and sequenced using the on DNBSEQ platform (DNBSEQ Technology). RNA-seq paired-end reads (read length 100 base pairs) were aligned to the mouse GRCm38 reference genome and read count for each gene was calculated using STAR Aligner (star2.7.6a) (55); Genes were compared for differential expression between the different treatments using edgeR (56) and were considered to be statistically expressed when the absolute fold change ≥2 and false discovery rate (FDR) <5%. These significantly expressed gene lists were subject to further functional annotation using enrichR (57-59) to identify altered pathways due to the corresponding treatments. For individual pathways, the Benjamini–Hochberg procedure was used to calculate FDR to adjust for multiple testing. RNA-seq data supporting the findings was deposited in BioProject, ID PRJNA848127.

**Quantitative RT-PCR**

Total RNA was isolated from transgenic mouse frozen tumor samples and coeliac ganglia with the RNAeasy kit (Qiagen) followed by RT-PCR with the thermoScript RT-PCR system (Invitrogen). Quantitative PCR was carried out with the Taqman probes Mm00627179 (murine *mycn*), JX952197 (human *MYCN*) and GAPDH Mm9999915.

**SUPPLEMENTARY TABLES**

**Supplementary Table S1**: list of cell lines screened for crizotinib, TAE-684, ceritinib alectinib and lorlatinib ALK inhibitors.

| **Cell Line** | ***MYCN*** | ***TP53*** | ***ALK*** | ***RAS/MAPK*** | ***ALK Status reference*** | **Crizotinib** (nM) | **TAE-684** (nM) | **Ceritinib** (nM) | **Alectinib** (nM) | **Lorlatinib** (nM) | **Immunoassay** |
| --- | --- | --- | --- | --- | --- | --- | --- | --- | --- | --- | --- |
|  | | | | | | Diagnosis | | | | | |
| CLB-Ge2 | Amp | WT | Amp; F1174V | WT | (12) | 188 ± 21 | 21.6 ± 3.6 | 59.7 ± 5.1 | 119.8 ± 21 | 30.7 ± 7.7 |  |
| IMR32 | Amp | WT | Partial amp | NF1, R69G | (14) | 934.3 ± 68.2 | 94.0 ± 32.4 | 862.6 ± 86.5 | 609.9 ± 20.65 | >10µM | yes |
| LAN5 | Amp | WT | R1275Q | WT | (51) | 258.4 ± 38.9 | 10.3 ± 0.3 | 170.1 ± 55.2 | 74.3 ± 11.9 | 81.4 ± 30.9 | yes |
| NB69 | Non-amp | WT | WT | WT | (51) | 634.3 ± 26.8 | 570.0 ± 36.7 | ND | 548 ± 31.8 | >10µM |  |
| NBLW | Amp | WT | R1275L | NF1, E91-* | (34) | 584.0 ± 41.6 | 38.5 ± 3.9 | 842.5 ± 91.5 | 488.5 ± 49.1 | >10µM |  |
|  | | | | | | Relapse/Post-treatment | | | | | |
| CHLA136 | Amp | WT | WT | WT | (60) | ND | 314.7 ± 36.3 | ND | >5µM | >10µM |  |
| CLB-BAR-Rec | Amp | WT | Amp; Del Ex4-11 | WT | (61) | 311.5 ± 28.4 | 3.4 ± 1.1 | 74.5 ± 10.1 | 151.7 ± 9.8 | 37.2 ± 11.7 |  |
| GIMEN | Non-amp | WT | WT | NF1, L551F | (14) | 1655.5 ± 320.5 | 500.0 ± 61.7 | 1594.7 ± 184.3 | >5µM | >10µM |  |
| LAN1 | Amp | Mut | F1174L | WT | (51); (50) | ND | 25.8 ± 9.6 | ND | 243.6 ± 17.4 | 3436.2 ± 1361.9 |  |
| LAN6 | Non-amp | WT | D1091N | KRAS, G12C | (50) | ND | 23.1 ± 3.8 | 567.8 ± 67.8 | 322.8 ± 55.1 | >10µM |  |
| NB-1* | Amp | WT | Amp; Del Ex2-3 | WT | (62) | 37.5 ± 5.9 | 3.3 ± 1.6 | 19.4 ± 4.1 | 11.7 ± 2.1 | 2.5 ± 0.3 |  |
| NB1691 | Amp | WT | Amplified | MAP2K2, V228M | This study | 857.5 ± 51.9 | 280.1 ± 81.3 | 916.4 ± 45.6 | 914.5 ± 59.7 | >10µM |  |
| NBLW-R | Amp | WT | F1174L | WT | (34) | 494.2 ± 43.5 | 30.2 ± 5.3 | 176.8 ± 28.4 | 307.3 ± 18.3 | 169.9 ± 20.6 | yes |
| NGP | Amp | WT | CNG | NF1, E1401* | (63) | 783.0 ± 157.5 | 179.3 ± 10.9 | 627.4 ± 78.9 | 609.1 ± 77.3 | >10µM |  |
| SHSY5Y | Non-amp | WT | F1174L | KRAS, G12V | (50) | 358.0± 49.5 | 57.4 ± 16.8 | 313 ± 77.7 | 249.5 ± 44.4 | 3674.8 ± 990.3 | yes |
| SKNAS | Non-amp | Mut | WT | NRAS, Q61K | (14) | 1230.3 ± 128.3 | 570.9 ± 91.2 | 1697.3 ± 249.7 | 2425.9 ± 142.2 | >10µM |  |
| SKnBe2C | Amp | Mut | CNG (sub pop) | NF1, Asn664* | (61) | 987.7 ± 13.71 | 119.8 ± 14.5 | 363.6 ± 60.4 | 665.5 ± 38.9 | >10µM |  |
| TR14 | Amp | WT | CNG | WT | (61) | ND | 97.8 ± 30.4 | ND | 186.3 ± 64.6 | >10µM |  |
|  | | | * Assume post treatment as established from patient who died from disseminated disease | | |  | | | ^1^Cell line sequencing from Newcastle Genetics Lab | | |
|  | | | WT, wild type; Mut, mutant; Del, deletion; ND, not determined; CNG, copy number gain | | |  | | |  | | |

**Supplementary Table S2:** Summary of the clinical and molecular characteristics of the five *ALK*-aberrant high-risk neuroblastoma (HR-NB) patient-derived xenografts. Model GR-NB4 was established within the MAPPYACTS program (49). We also indicate whether each PDX was used for *ex vivo* high throughput drug screenings or *in vivo* efficacy of ALK inhibitors. ALKi: ALK inhibitors; VAF: variant allele fraction; ALT: alternative lengthening of telomeres; NA: not applicable; WT: wild type, Del: deletion; Not avail.: not available.

| *ALK*-aberrant HR-NB  PDX MODELS | CLINICAL DATA | | | | SOMATIC GENETIC ALTERATIONS | | | | | | | EXPERIMENT | |
| --- | --- | --- | --- | --- | --- | --- | --- | --- | --- | --- | --- | --- | --- |
|  | **PDX timepoint** | **Stage** | **Gender** | **Previous ALKi** | ***MYCN*** | ***ALK*** (VAF) | ***TP53*** | ***ATRX*** (VAF) | **Copy number alteration** | **ALT** | **Other relevant somatic genetic abnormalities** | ***Ex***  ***vivo*** | ***In***  ***vivo*** |
| GR-NB4 | Relapse | 4 | F | No | A | A | WT | WT | Del. 5p, 6p, 9q; focal gain 5pq | NA | CDKN2A/B homo del | Yes | Yes |
| IC-pPDX-75 | Relapse | 4 | F | Crizotinib | WT | F1174L (65%) | WT | c.6391C>T (34%) | Del. 1q, 11q; gain 5p, 7q, 17q | Not avail. |  | Yes | No |
| IC-pPDX-112 | Diagnosis | 4 | M | No | A | Intron 3 of ALK Amp | WT | WT | Del. 1p, 3p, 4p; gain 2p, 17q | NA | High mutational load | Yes | Yes |
| HSJD-NB-011 | Relapse | 4 | M | No | A | I1171N (80%) | WT | WT | Not avail. | Negative |  | Yes | No |
| HSJD-NB-012 | Relapse | 4 | M | No | A | F1174C (30%) | WT | WT | Del. 1p, 7q, 11q, 17p; gain 1q, 17q | Negative |  | Yes | Yes |

**Supplementary Table S3:** ALK, Ki67, P53 and cleaved caspase 3 immunohistochemical scoring of PDX tumors taken from the 3-day treatment cohort, according to score by two independent pathologists (H&E, Haematoxylin and Eosin staining).

|  |  | **H&E** | **ALK membrane &**  **cytoplasm** | | | **Ki67 nucleus** | | | **P53 nucleus** | | | **P21 nucleus** | | | **Cleaved Caspase 3 nucleus** | | |
| --- | --- | --- | --- | --- | --- | --- | --- | --- | --- | --- | --- | --- | --- | --- | --- | --- | --- |
| **PDX** | **Treatment** | **Tumor Surface** | **%** | **Intensity** | **Score Histo** | **%** | **Intensity** | **Score Histo** | **%** | **Intensity** | **Score Histo** | **%** | **Intensity** | **Score Histo** | **%** | **Intensity** | **Score Histo** |
| GR-NB4 | Vehicle | 90 | 98 | 3 | 3 | 87 | 3 | 3 | 82 | 2 | 2 | 25 | 3 | 1 | 2 | 3 | 0 |
| GR-NB4 | CAV | 90 | 98 | 3 | 3 | 90 | 3 | 3 | 88 | 2 | 2 | 27 | 3 | 1 | 5 | 3 | 0 |
| GR-NB4 | CAV &  lorlatinib | 90 | 98 | 3 | 3 | 83 | 3 | 3 | 85 | 2 | 2 | 25 | 3 | 1 | 8 | 3 | 0 |
| GR-NB4 | Lorlatinib | 90 | 94 | 3 | 3 | 72 | 3 | 2 | 89 | 2 | 1 | 17 | 3 | 0 | 4 | 3 | 0 |
| GR-NB4 | Lorlatinib & Idasanutlin | 90 | 98 | 3 | 3 | 73 | 3 | 2 | 80 | 2 | 2 | 22 | 3 | 1 | 14 | 3 | 0 |
| GR-NB4 | Idasanutlin | 90 | 98 | 3 | 3 | 77 | 3 | 2 | 94 | 3 | 2 | 22 | 3 | 1 | 4 | 3 | 0 |
| IC-pPDX-112 | Vehicle | 90 | 98 | 2 | 2 | 63 | 3 | 2 | 4 | 1 | 0 | 75 | 3 | 2 | 1 | 3 | 0 |
| IC-pPDX-112 | CAV | 37 | 93 | 2 | 2 | 58 | 3 | 2 | 70 | 1 | 0 | 57 | 3 | 2 | 1 | 3 | 0 |
| IC-pPDX-112 | CAV &  Lorlatinib | 72 | 91 | 2 | 2 | 64 | 3 | 2 | 37 | 2 | 0 | 69 | 2 | 2 | 0 | 3 | 0 |
| IC-pPDX-112 | Lorlatinib | 84 | 93 | 2 | 2 | 65 | 3 | 2 | 44 | 2 | 1 | 68 | 3 | 2 | 1 | 3 | 0 |
| IC-pPDX-112 | Lorlatinib & Idasanutlin | 90 | 90 | 1 | 1 | 72 | 3 | 2 | 40 | 2 | 1 | 55 | 3 | 2 | 0 | 3 | 0 |
| IC-pPDX-112 | Idasanutlin | 63 | 60 | 1 | 1 | 62 | 3 | 2 | 65 | 2 | 1 | 57 | 3 | 2 | 1 | 3 | 0 |
| HSJD-NB-012 | Vehicle | 90 | 57 | 1 | 1 | 78 | 3 | 2 | 63 | 1 | 1 | 5 | 2 | 0 | 4 | 3 | 0 |
| HSJD-NB-012 | CAV | 90 | 47 | 1 | 0 | 80 | 3 | 2 | 51 | 1 | 1 | 11 | 3 | 0 | 2 | 3 | 0 |
| HSJD-NB-012 | CAV &  lorlatinib | 90 | 57 | 1 | 1 | 80 | 3 | 2 | 74 | 1 | 1 | 1 | 1 | 0 | 8 | 3 | 0 |
| HSJD-NB-012 | Lorlatinib | 68 | 73 | 2 | 1 | 73 | 3 | 2 | 65 | 1 | 1 | 1 | 1 | 0 | 10 | 3 | 0 |
| HSJD-NB-012 | Lorlatinib & Idasanutlin | 90 | 73 | 1 | 1 | 77 | 3 | 2 | 54 | 2 | 1 | 9 | 3 | 0 | 8 | 3 | 0 |
| HSJD-NB-012 | Idasanutlin | 90 | 18 | 1 | 0 | 79 | 3 | 2 | 63 | 2 | 1 | 10 | 2 | 0 | 5 | 3 | 0 |
| One-way ANOVA |  |  | P=0.0003*** |  |  | P=0.0002*** |  |  | P=0.0006*** |  |  | P<0.0001**** |  |  | P=0.0091** |  |  |

**Supplementary Table S4:** GR-NB4 PDX Relative Tumor Volumes (up to day 10) in the Lorlatinib-Idasanutlin combination study.

|  | **Day** | | | |
| --- | --- | --- | --- | --- |
| **Study Arm** | **1** | **4** | **7** | **10** |
| **Vehicle** | 1.00 | 1.10 | 2.74 | 4.10 |
|  | 1.00 | 1.00 | 1.17 | 2.37 |
|  | 1.00 | 1.00 | 1.19 | 1.89 |
|  | 1.00 | 1.00 | 1.17 | 1.17 |
|  | 1.00 | 1.56 | 1.94 | 2.79 |
| **Idasanutlin** | 1.00 | 2.02 | 3.14 | 3.14 |
|  | 1.00 | 0.58 | 0.58 | 0.58 |
|  | 1.00 | 0.92 | 0.83 | 1.89 |
|  | 1.00 | 1.11 | 1.68 | 1.68 |
|  | 1.00 | 1.56 | 2.29 | 2.29 |
| **Lorlatinib** | 1.00 | 0.51 | 0.64 | 1.00 |
|  | 1.00 | 0.53 | 0.53 | 0.53 |
|  | 1.00 | 1.09 | 1.51 | 1.51 |
|  | 1.00 | 1.00 | 1.00 | 1.20 |
|  | 1.00 | 1.00 | 1.00 | 1.00 |
| **Lorlatinib & Idasanutlin** | 1.00 | 0.22 | 0.22 | 0.22 |
|  | 1.00 | 0.58 | 0.30 | 0.30 |
|  | 1.00 | 0.61 | 0.69 | 0.69 |
|  | 1.00 | 0.36 | 0.36 | 0.36 |
|  | 1.00 | 0.51 | 0.64 | 0.64 |
|  | 1.00 | 0.63 | 0.63 | 0.63 |

**SUPPLEMENTARY FIGURES**

**Supplementary Figure S1**: (A) High density SNP array (Illumina 850K) of NB1691 cell line showing *MYCN*, *ALK* and *MDM2* amplification analyzed using Nexus software. (B) Array CGH Copy number profile of IC-pPDX-112 model carried an intron 3 *ALK* amplification. B In the chr2 p gained region, *SOX11, MYCN* and the intron 3 of *ALK* (chr2:29800001-29900000) are amplified.

**Supplementary Figure S2**: (A) Correlation between independent high-throughput drug screenings on PDX-derived tumor cells (two different experiments, same PDX model, different passages). The Pearson’s correlation coefficient (*r*) is indicated. R1: replicate 1; R2: replicate 2. (B) Distribution of variability for each PDTC model in control wells. Data normalized by DMSO within each plate.

**Supplementary Figure S3:** (A) 3-Day treatment of Th-*MYCN* tumor-bearing animals with ALK inhibitor panel. (B) *Ex vivo* analysis of pY1586 ALK / total ALK expression using immunoassay of tumor lysates from the lorlatinib survival study in the Th-*ALK*^F1174L^/*MYCN* GEMM. Unpaired t-test vehicle versus lorlatinib p=0.0018. (C) qRT-PCR of human *MYCN* or murine *mycn* in tumor samples from the lorlatinib survival study in the Th-*ALK*^F1174L^/*MYCN* GEMM. Unpaired t test vehicle versus lorlatinib (Th-*ALK^F1174L^/MYCN*) p<0.0001. (D) Classification of “Poor Responders” and “Early Responders” from the lorlatinib survival study in the Th-*ALK*^F1174L^/*MYCN* GEMM, according to the volumetric response at day 7, as measured by MR imaging. Graph shows % tumor volume change between day 1 and day 7, with each bar representing an individual animal. (E) Heatmap of ALK and adrenergic signature gene expression taken following analysis of RNA sequencing of tumor samples from the end of the lorlatinib survival study.

**Supplementary Figure S4:** (A) Heat map for the 363 uniquely differentially expressed genes between vehicle and CAV-lorlatinib combination treated Th-*ALK^F1174L^/MYCN* tumors. (B) (i) Hierarchical clustering and heatmap representation of the Lambertz’s ALK-genset 76 gene expression across the 14 tumor samples. For all heat maps, green denotes the standardized z-score>0, a higher gene expression; and blue denotes the standardized z-score<0, a lower gene expression compared to other samples. (ii) log2 trimmed mean of M values (TMM) for *Etv5* expression. (C) Median gene rank of the NCC-Boeva-like gene signature versus the median gene rank of the noradrenergic-Boeva signature.

**Supplementary Figure S5**: (A) Heatmap of expression of 35 selected genes across all the treatment groups. ADRN (Noradrenergic) and MES-like (Mesenchymal-like). (B) (i) Immunoblot of 3 the vehicle tumors from each PDX model, taken at the end of the survival experiment. (ii) Immunoassay for total ALK (average of 2 technical repeats) One-way ANOVA p=0.0264*. (C) Densitometry (normalized to GAPDH) of immunoblots shown in Figure Bii, Cii and Eii: (C.i) GR-NB4, one-way ANOVA p=0.0051** (pAKT/AKT), one-way ANOVA p=0.0037** (pERK/ERK); (C.ii) PDX112, one-way ANOVA p=0.0017** (pERK/ERK); (C.iii) HSJD102. Mean of 2 technical repeats. (D) ALK expression distribution evaluated by RNA sequencing of clinical neuroblastomas (NB), in comparison to Anaplastic Large Cell Lymphoma (ALCL) and other pediatric tumors (other). Wilcoxon test FDR-adjusted p value: NB ALK mutation versus other NB p=0.0515**; NBL ALK mutation versus ALCL p=0.0586*; NB ALK mutation versus other p=0.00016***. (TPM: transcript count per million).

**Supplementary Figure S6:** (A.i-iv.) *In vitro* cell growth curves corresponding to Figure 5A. (B.i-iv.) *In vitro* cell growth curves corresponding to Figure 5B. (C) GR-NB4 PDX lorlatinib and idasanutlin combination experiment, in which animals treated with the combination were left after treatment had finished until tumors had relapsed and achieved ethical size limit.

SUPPLEMENTARY REFERENCES

37.	Yadav B, Pemovska T, Szwajda A, Kulesskiy E, Kontro M, Karjalainen R, et al. Quantitative scoring of differential drug sensitivity for individually optimized anticancer therapies. Sci Rep. 2014;4:5193.

38.	Kilkenny C, Browne WJ, Cuthill IC, Emerson M, Altman DG. Improving bioscience research reporting: The ARRIVE guidelines for reporting animal research. J Pharmacol Pharmacother. 2010;1(2):94-9.

39.	Workman P, Aboagye EO, Balkwill F, Balmain A, Bruder G, Chaplin DJ, et al. Guidelines for the welfare and use of animals in cancer research. Br J Cancer. 2010;102(11):1555-77.

40.	Jamin Y, Tucker ER, Poon E, Popov S, Vaughan L, Boult JK, et al. Evaluation of clinically translatable MR imaging biomarkers of therapeutic response in the TH-MYCN transgenic mouse model of neuroblastoma. Radiology. 2013;266(1):130-40.

41.	Tucker ER, Tall JR, Danielson LS, Gowan S, Jamin Y, Robinson SP, et al. Immunoassays for the quantification of ALK and phosphorylated ALK support the evaluation of on-target ALK inhibitors in neuroblastoma. Mol Oncol. 2017;11(8):996-1006.

55.	Dobin A, Davis CA, Schlesinger F, Drenkow J, Zaleski C, Jha S, et al. STAR: ultrafast universal RNA-seq aligner. Bioinformatics. 2013;29(1):15-21.

56.	Robinson MD, McCarthy DJ, Smyth GK. edgeR: a Bioconductor package for differential expression analysis of digital gene expression data. Bioinformatics. 2010;26(1):139-40.

57.	Chen EY, Tan CM, Kou Y, Duan Q, Wang Z, Meirelles GV, et al. Enrichr: interactive and collaborative HTML5 gene list enrichment analysis tool. BMC Bioinformatics. 2013;14:128.

58.	Kuleshov MV, Jones MR, Rouillard AD, Fernandez NF, Duan Q, Wang Z, et al. Enrichr: a comprehensive gene set enrichment analysis web server 2016 update. Nucleic Acids Res. 2016;44(W1):W90-7.

59.	Xie Z, Bailey A, Kuleshov MV, Clarke DJB, Evangelista JE, Jenkins SL, et al. Gene Set Knowledge Discovery with Enrichr. Curr Protoc. 2021;1(3):e90.

60.	Koneru B, Lopez G, Farooqi A, Conkrite KL, Nguyen TH, Macha SJ, et al. Telomere Maintenance Mechanisms Define Clinical Outcome in High-Risk Neuroblastoma. Cancer Res. 2020;80(12):2663-75.

62.	Okubo J, Takita J, Chen Y, Oki K, Nishimura R, Kato M, et al. Aberrant activation of ALK kinase by a novel truncated form ALK protein in neuroblastoma. Oncogene. 2012;31(44):4667-76.
