## Supplementary figures and images for "COMBINATION THERAPIES TARGETING ALK-ABERRANT NEUROBLASTOMA IN PRECLINICAL MODELS"

### Figure S1

#### Supplementary Figure S1

A

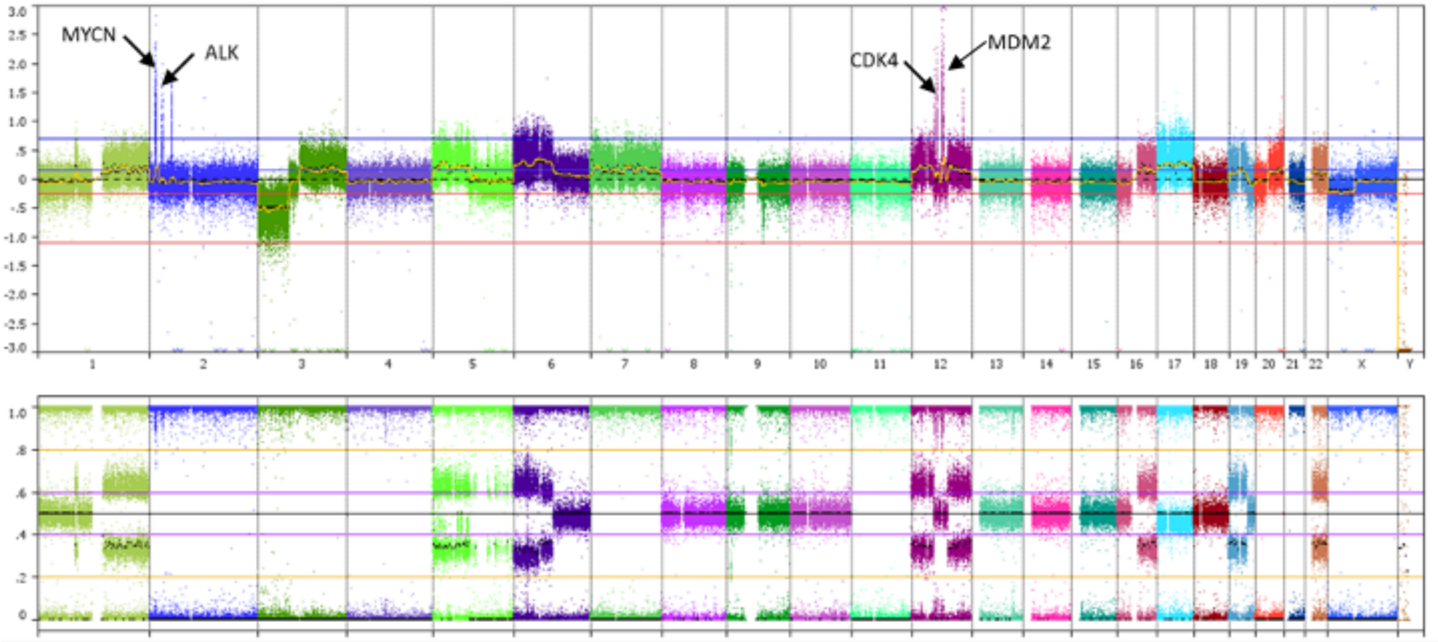


B

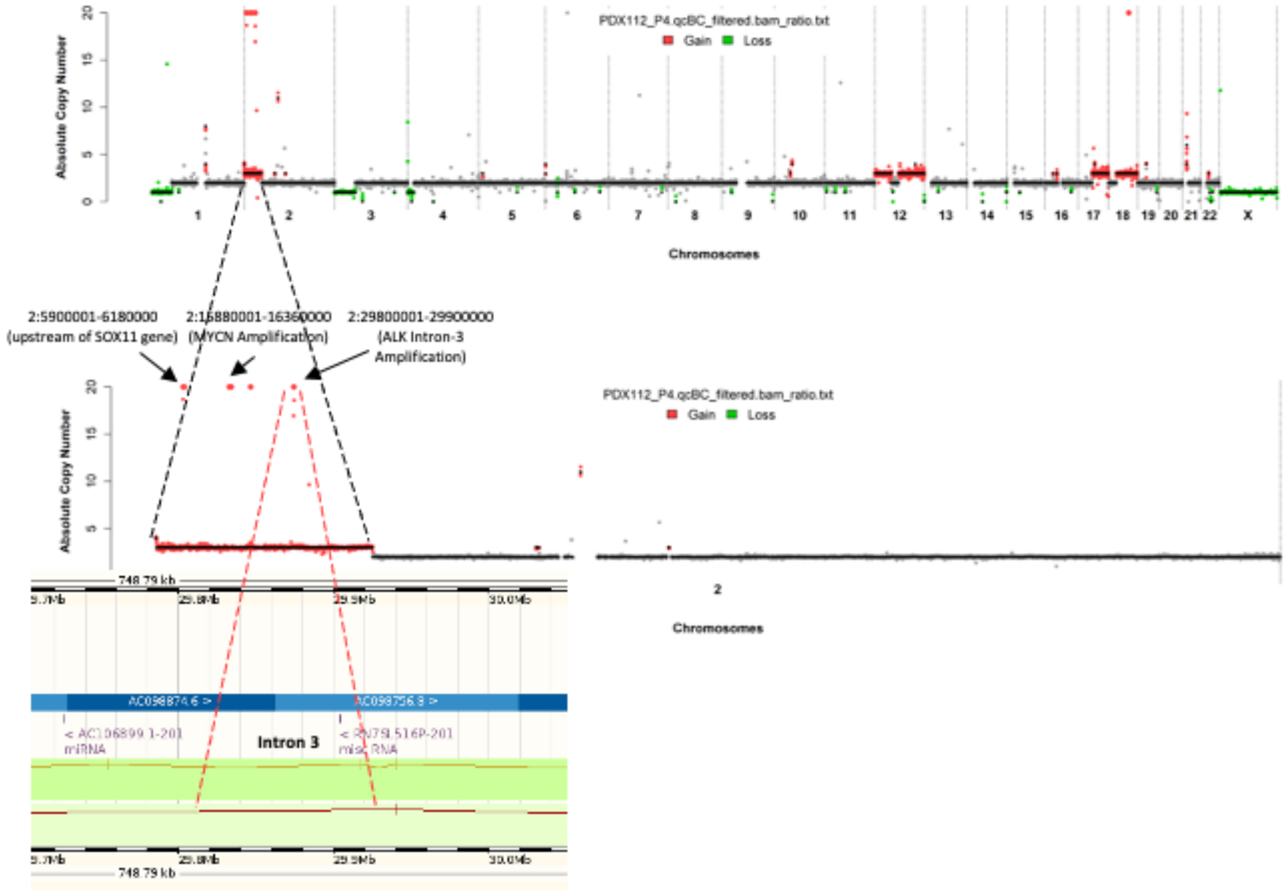


**Amplified ALK region : chr2:29800001-29900000**

### Figure S2

#### Supplementary Figure S2

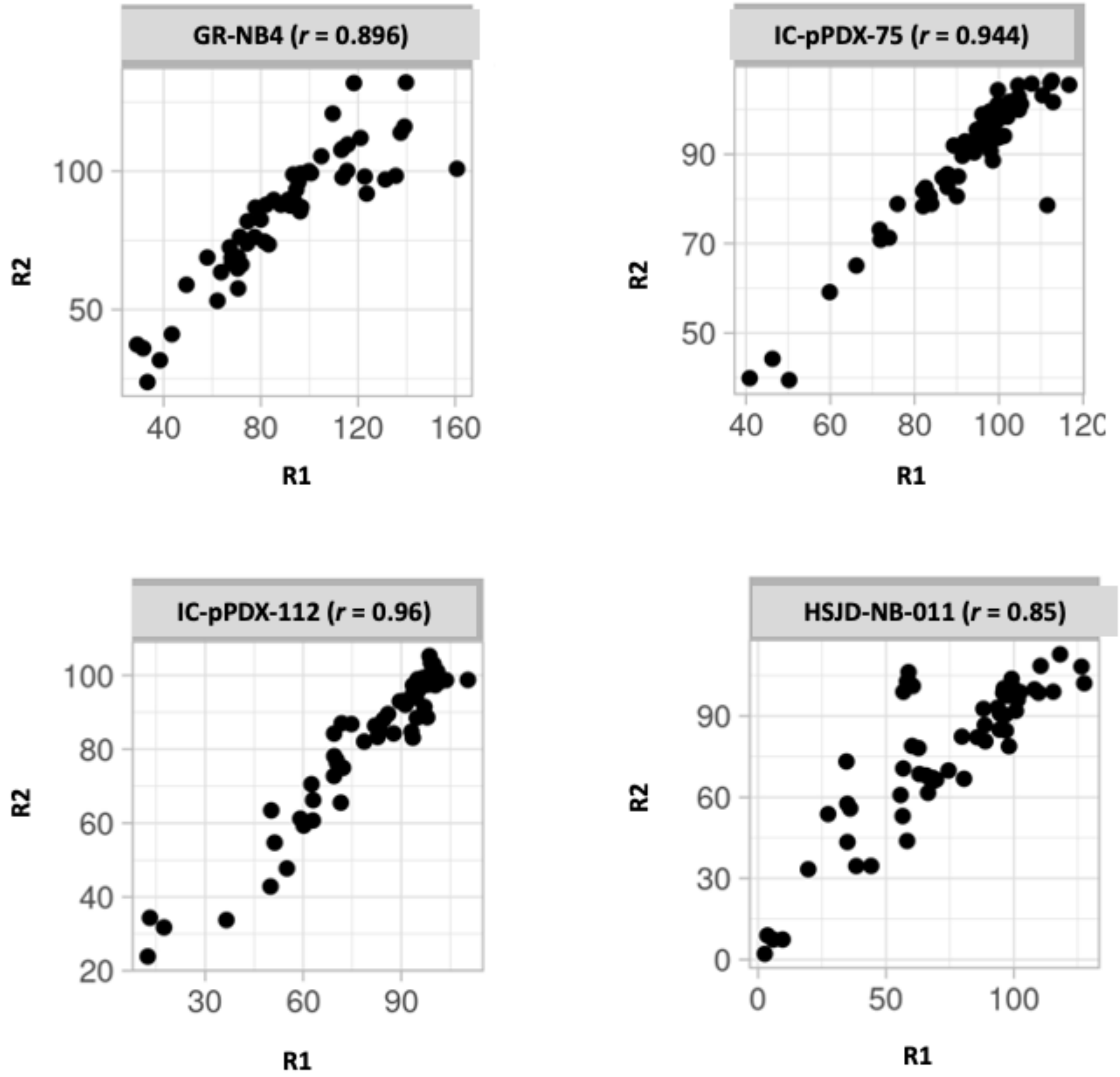

### Figure S3

#### Supplementary Figure S3

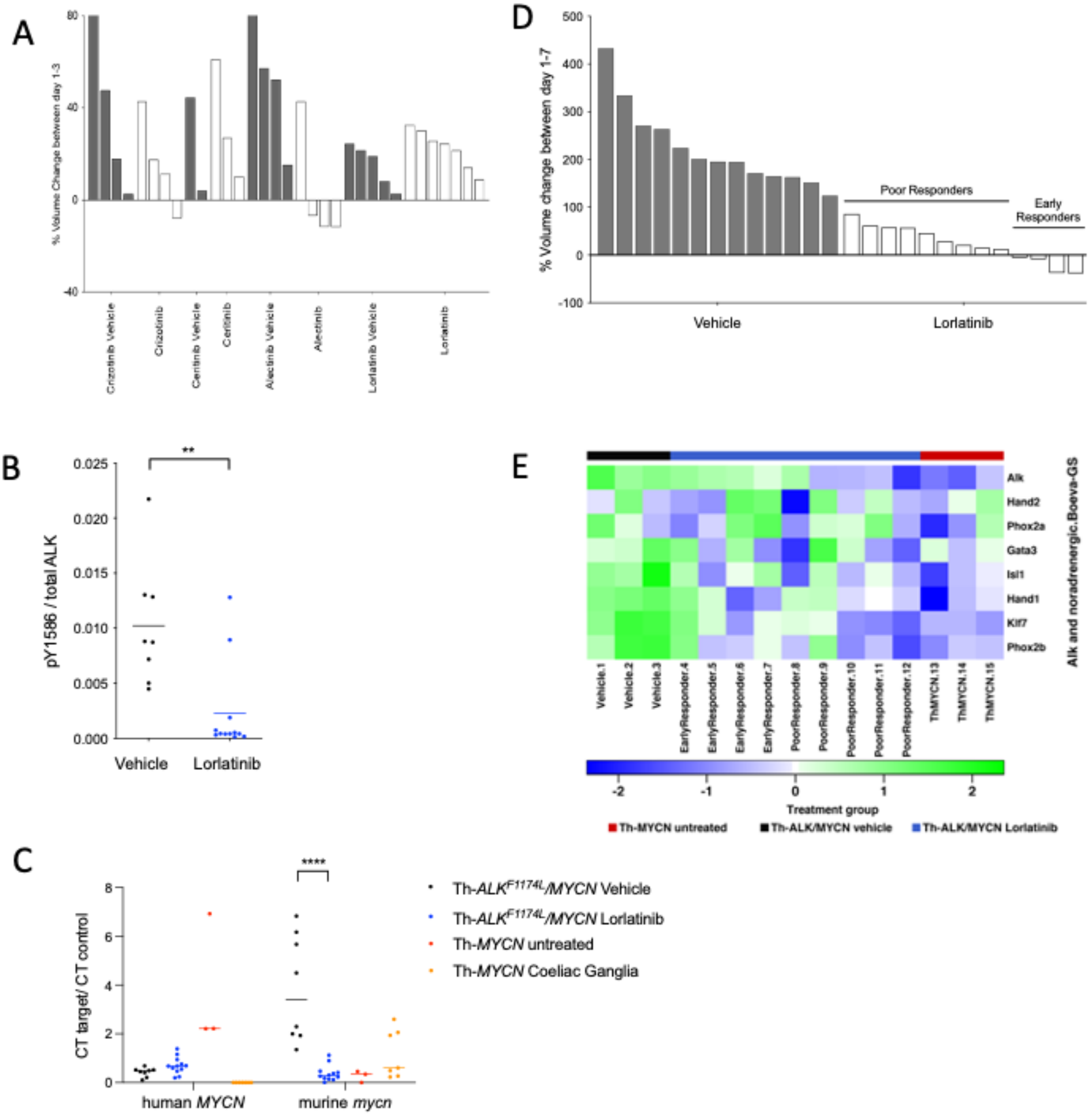

### Figure S4

#### Supplementary Figure S4

A.

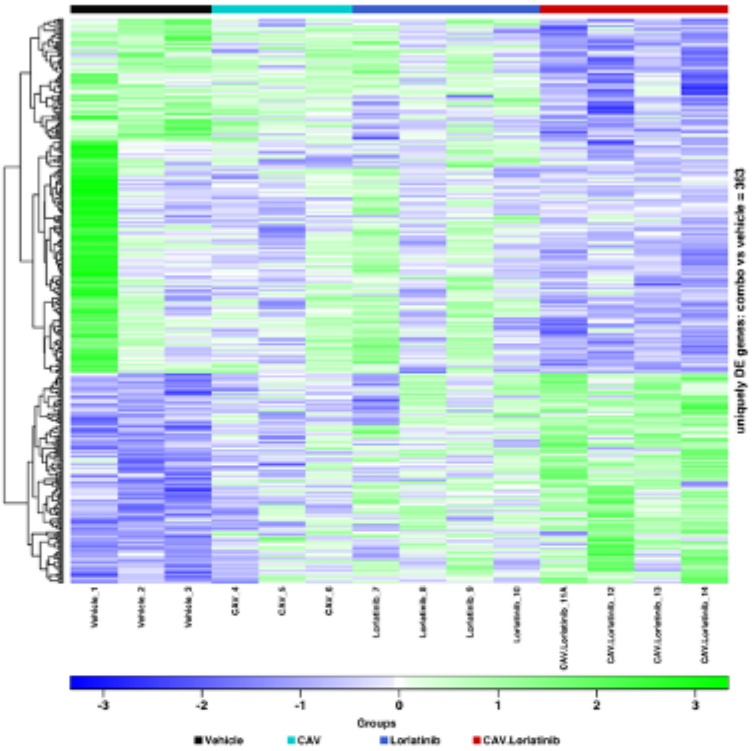


C.

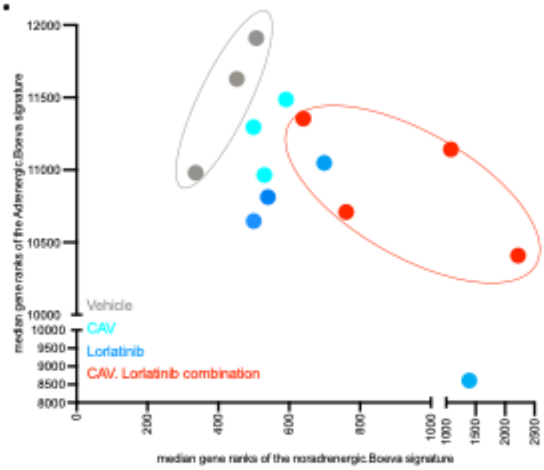


B.i.

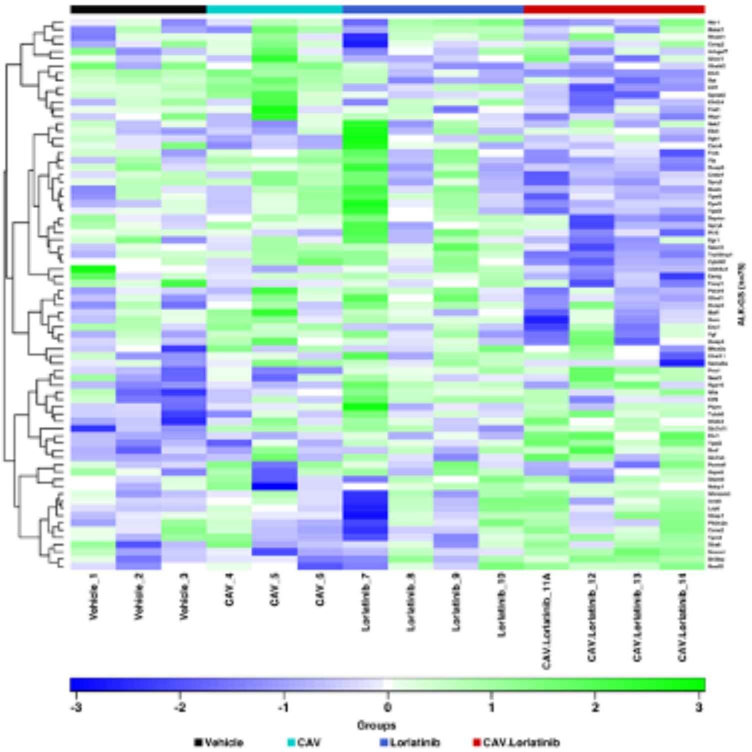


ii.

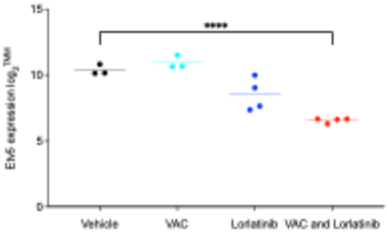

### Figure S5

#### Supplementary Figure S5

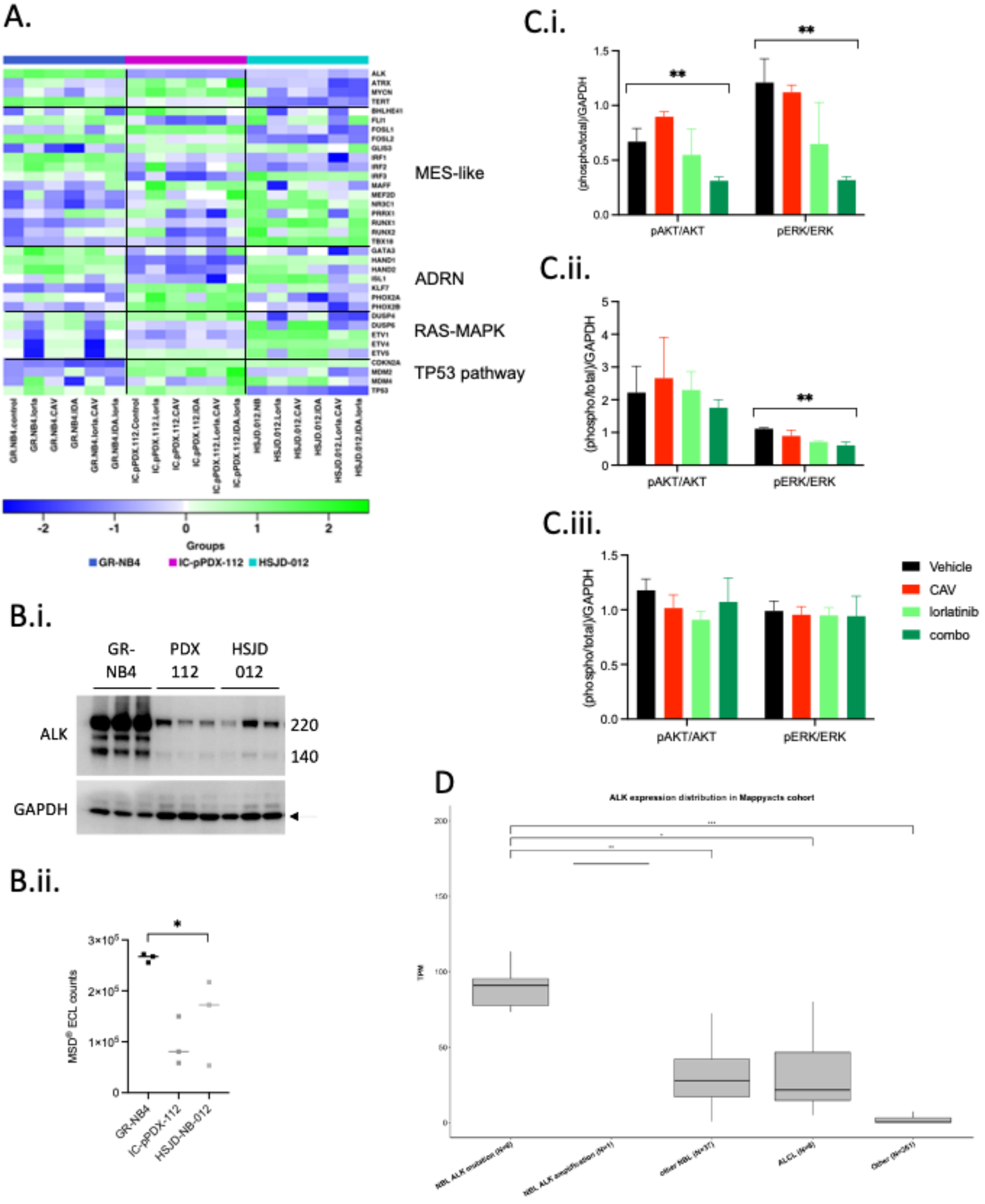

### Figure S6

#### Supplementary Figure S6

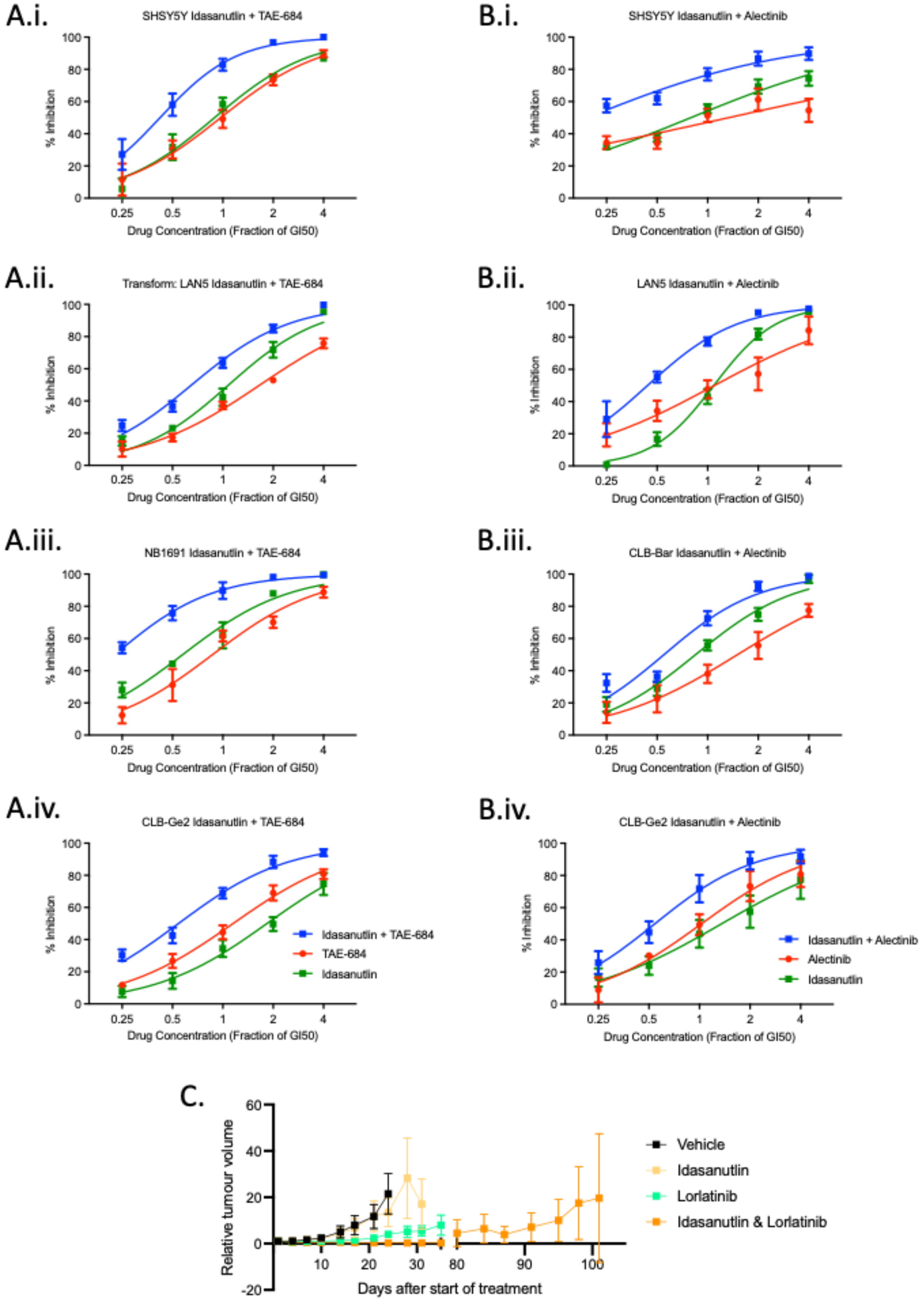
